# Exploring polycyclic scaffolds as adamantane replacements in M2 channel inhibitors of Influenza A virus

**DOI:** 10.1101/2025.07.26.666854

**Authors:** Andreea L. Turcu, Rosana Leiva, Chunlong Ma, Kyriakos Georgiou, José M. Brea, M. Isabel Loza, Cristina Val, Lieve Naesens, Jun Wang, Antonios Kolocouris, Santiago Vázquez

## Abstract

The increasing resistance of influenza A viruses to adamantane-based antivirals underscores the need for new inhibitors targeting both wild-type (WT) and mutant M2 ion channels. Here, we report the synthesis and biological evaluation of polycyclic cage amines designed to replace the adamantane scaffold as M2 inhibitors. These include ring-contracted and ring-expanded analogues, evaluated both as primary amines and as aryl-/heteroaryl-substituted derivatives.

Most of the polycyclic amines inhibited the WT M2 channel as demonstrated by electrophysiological assays. Among them, compound **10**, a 3,4,8,9-tetramethyltetracyclo[4.4.0.0³.⁶.0⁴.⁸]decan-1-amine, emerged as a triple blocker active against M2 WT, M2 L27F, and M2 V27A channels. In contrast, compound **6c**, a noradamantane–isoxazole derivative, showed selective inhibition of the S31N mutant. Although no antiviral activity was observed against influenza A virus in infected cell assays, both compounds **6c** and **10** displayed significant antiviral activity against human coronavirus 229E. Furthermore, compound **10** demonstrated favourable pharmacokinetic properties.

MD simulations show that noradamantane **6c** binds inside the M2 S31N pore, with its ammonium forming H-bonds to Asn31 and the isoxazole positioned near Val27, restricting water entry. In contrast, larger polycyclic amines likely cannot access the pore due to steric hindrance.

## Introduction

Influenza A virus remains a major global health challenge, causing significant morbidity and mortality through seasonal epidemics and five documented pandemics since 1889, including the 1918 Spanish flu and the 2009 H1N1 pandemic. ^1^ Although vaccines are the primary strategy for influenza prevention, their effectiveness is limited by the rapid antigenic drift and shift of the virus, which necessitate constant reformulation and prediction of circulating strains.^2–3^ These limitations highlight the critical need for effective antiviral drugs that can serve both as therapeutic agents and as a first line of defense during pandemic emergence.

For decades, treatment of influenza virus infections relied on M2 ion channel inhibitors, particularly those based on the adamantane scaffold, such as amantadine and rimantadine. ^4^ These agents function by blocking the proton channel activity of the M2 protein, thereby inhibiting viral uncoating and subsequent replication.^4–5^ However, these drugs are no longer recommended for clinical use due to increase rate of resistant to the adamantanes.^6,7^ Resistance-associated mutations primarily occur in the transmembrane domain of the M2 protein, with the S31N substitution present in the vast majority of resistant strain. ^8–11^ In addition, other mutations such as V27A and L26F have been identified in circulating viruses, and in some cases, double mutants, such as V27A/S31N or L26I/S31N have also been detected.^11–13^ Although these variants remain less prevalent, their potential to expand under antiviral selective pressure or emerge from zoonotic reservoirs raises concern. Furthermore, while neuraminidase inhibitors such as oseltamivir and zanamivir have served as alternative therapies, resistance to these agents has also been reported.^14,15^ This rising resistance levels call for the design of next-generation antivirals active against resistant influenza A strains.

In response to this challenge, the researchers have developed a variety of adamantane-based antiviral compounds aimed at overcoming resistance. ^16–30^ One key strategy involved attaching aryl or heteroaryl rings to the amantadine amino group via a methylene bridge, allowing retention of the amine’s basicity. These second-generation inhibitors have been extensively studied through structure–activity relationship (SAR) analyses, focusing on modifications to both the adamantyl and aryl groups to enhance antiviral activity.^16–20^ This work has led to the identification of several potent inhibitors against the M2 S31N mutant virus, including compounds **1**,^16^ **2**^17^**, 3**^18^ (Figure 1). Among these compounds, phenol **1**, bearing a 2-hydroxy-4-methoxyphenyl group, and compound **3**, featuring a 2-bromothiophenyl group, acted as dual blockers of M2 WT and S31N channels by electrophysiological (EP) assays and inhibited replication of viral strains bearing these channels *in vitro*. In contrast, compound **2** which contains a 5-(2-thiophenyl)isoxazolyl moiety, selectively blocked the M2 S31N channel and inhibited only the corresponding virus *in vitro*. Additionally, some of these compounds have been further optimized by introducing a 3-hydroxy group to the adamantane core to improve their pharmacokinetic (ADME) properties ^19–20^.

**Figure 1.**
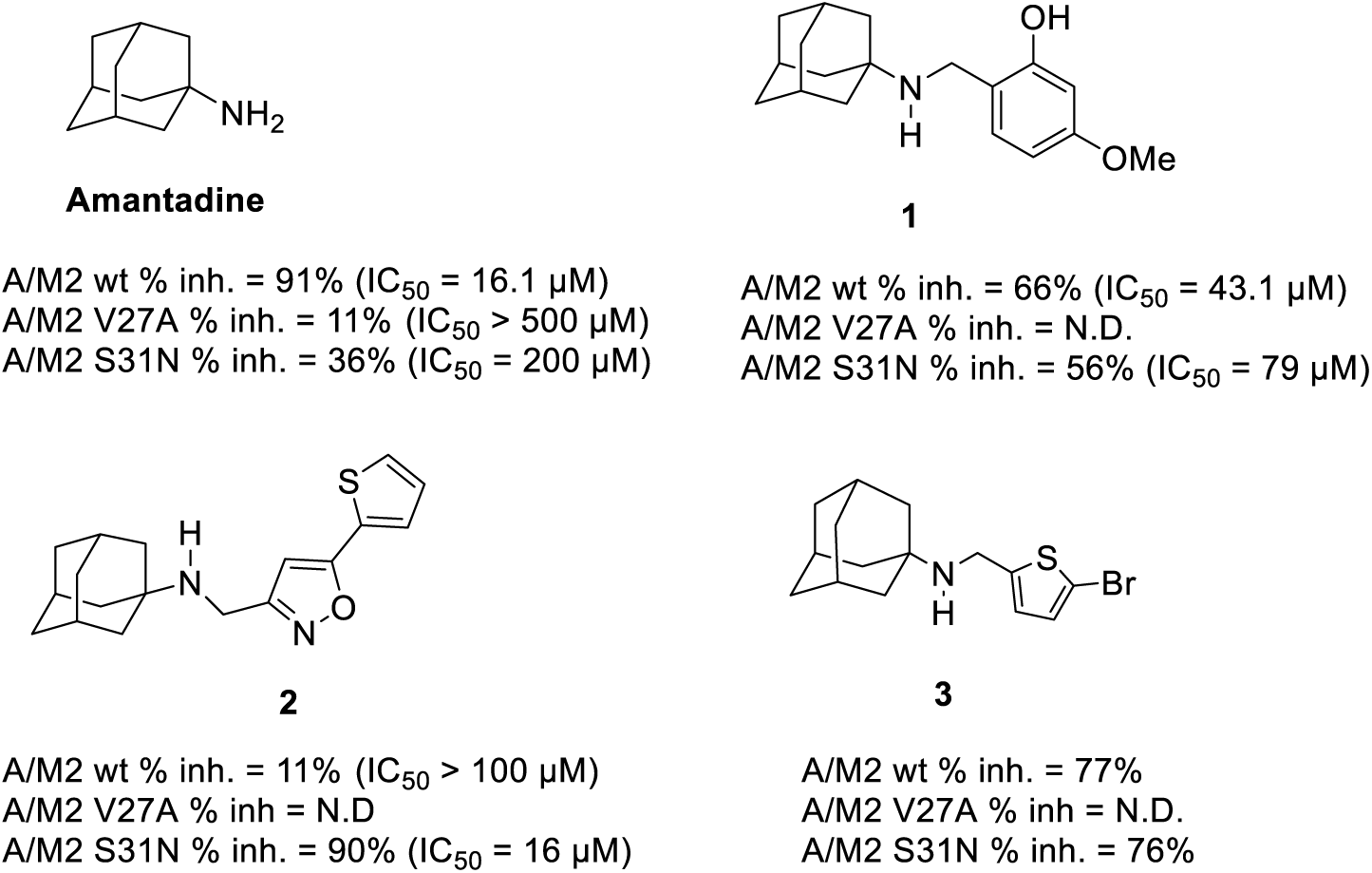
Structures of amantadine and some reported derivatives with potent activities against WT and/or mutant A/M2 channels. The inhibition percentages (determined at 100 μM for two minutes) and the IC_50_ values shown are previously reported values determined by TEVC assay.^16–18,25^ N.D. = Not Determined.

Computational and structural studies have shown that compound **2** binds within the channel pore of the M2 protein, specifically interacting with residues V27 to H37 in the S31N mutant, and V27 to G34 in the WT channel. Importantly, the binding orientation differs between the two forms: in M2-WT, the compound binds with its aromatic group facing the *C*-terminus, while in M2-S31N, the aromatic group is oriented toward the *N*-terminus. ^17,18, 31–32^ The observed flip-flop binding emphasizes the structural divergence of WT and mutant channels and the need for derivatives that block both to achieve broad-spectrum activity.

Thus, in previous work, we replaced the amantadine core in compounds **1–3** with 16 structurally diverse analogs, including 2-alkyl-2-adamantylamines, 3-substituted amantadine, rimantadine and its dialkyl adducts in the bridgehead carbon atom, 4-(1-adamantyl)benzenamine, diamantylamine or the triamantylamine, *tert*-alkyl amines, and a polycyclic cage amine. We observed that blocking M2 S31N is sterically constrained, as even a minimal extension, such as inserting a single carbon between the adamantyl and amino groups (e.g., using rimantadine instead of amantadine), abolished activity against this mutant. Furthermore, although some of the analogues exhibited moderate antiviral potency (∼20 μM) against viruses carrying the S31N M2 channel, they did not block M2 S31N in EP assays, underlying an additional mechanism of antiviral activity. Moreover, their low metabolic stability further underscores the need for structural optimization.^29^

Building on these findings, we sought to expand the chemical space of M2 ion channel blockers by systematically replacing the adamantane scaffold with a variety of polycyclic cage amines of different sizes and shapes. Our goal was to assess their ability, either as standalone cores or as aryl/heteroaryl derivatives (as in compounds **1–3**, Figure 1), to inhibit the WT M2 channel and drug-resistant variants such as L26F, V27A, and S31N, as determined by EP. This approach aimed to provide insight into the steric and spatial constraints of the M2 binding site and how mutations affect ligand accommodation.

The set of polycyclic scaffolds included previously reported structures such as oxadamantane (**4, 5**); ring-contracted structures such as noradamantane (**6**), tricyclo[3.3.1.0³,⁷]nonane (**7**), and tricyclo[3.3.0.0³,⁷]octane (**8, 9**); as well as larger-volume scaffolds, including tetracyclo[4.4.0.0³.⁶.0⁴.⁸]decane (**10**) and 4-azatetracyclo[5.4.2.0².⁶.0⁸^.11^]tridecane (**11**) (Figure 2).

**Figure 2.**
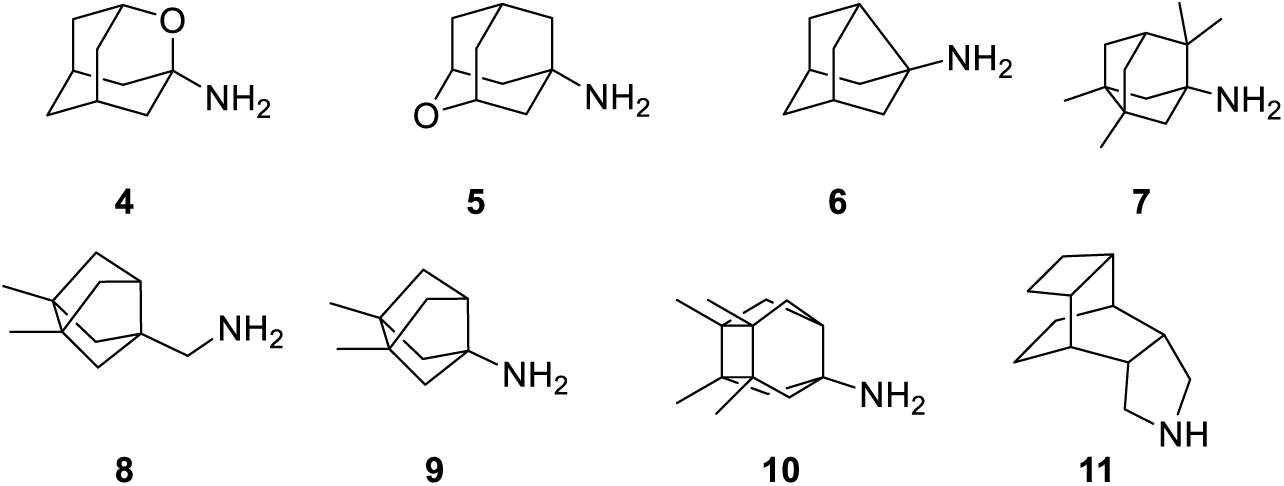
Polycyclic cage amines **4–11**, including both previously reported inhibitors of influenza A M2 channels (compounds **4**, **5**, **6**, **8**, **9**, **11**) and new scaffolds (compounds **7** and **10**) (see Table 1).

**Table 1.**
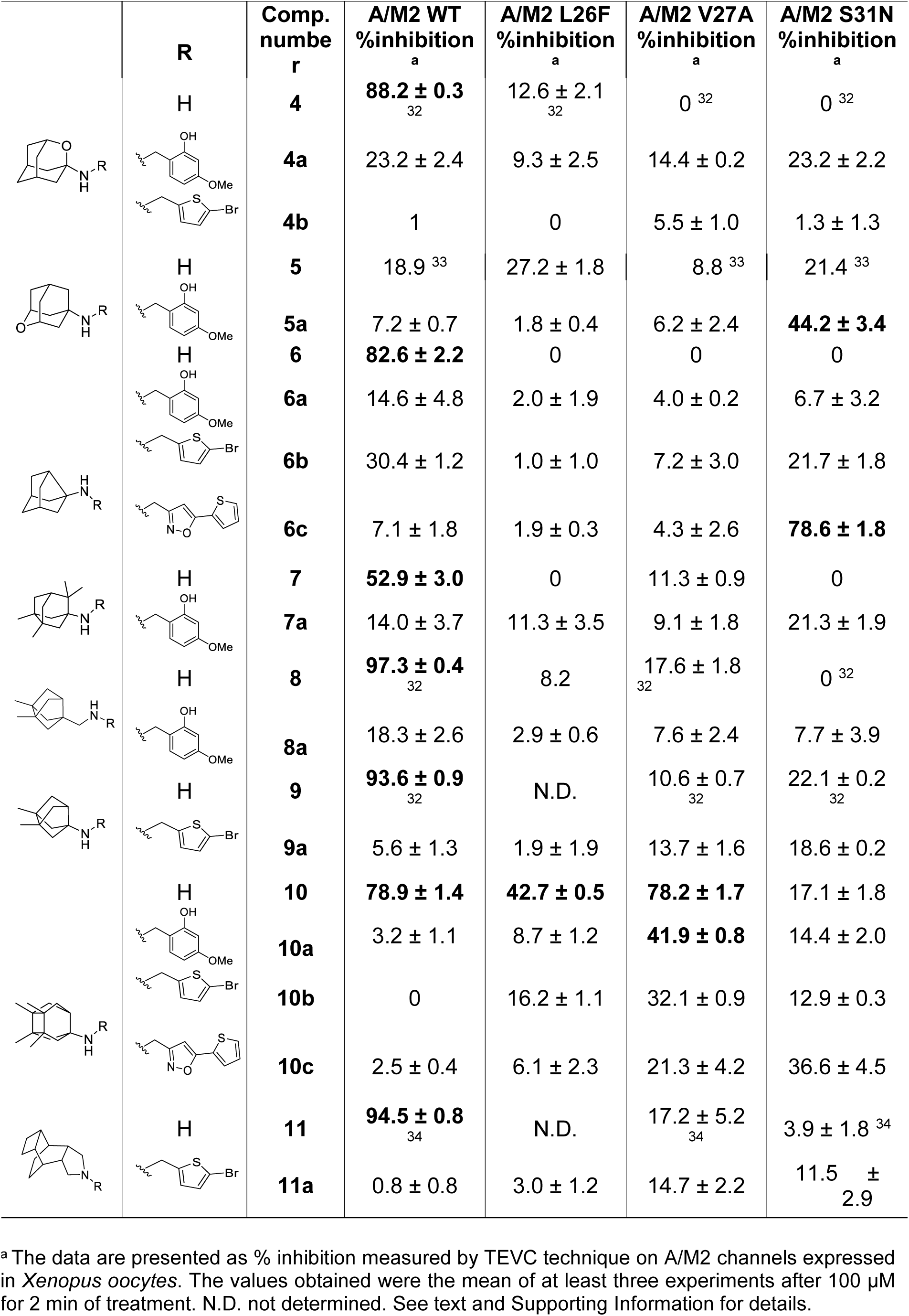
Structure and inhibitory effect of the new compounds on A/M2 WT, L26F, V27A and S31N proton channels functions using TEVC assay.

|  | R | Comp. number | A/M2 WT %inhibition<sup>a</sup> | A/M2 L26F %inhibition<sup>a</sup> | A/M2 V27A %inhibition<sup>a</sup> | A/M2 S31N %inhibition<sup>a</sup> |
| --- | --- | --- | --- | --- | --- | --- |
|  | H | **4** | **88.2 ± 0.3**<sub>32</sub> | 12.6 ± 2.1<sub>32</sub> | 0<sub>32</sub> | 0<sub>32</sub> |
|  |  | **4a** | 23.2 ± 2.4 | 9.3 ± 2.5 | 14.4 ± 0.2 | 23.2 ± 2.2 |
|  |  | **4b** | 1 | 0 | 5.5 ± 1.0 | 1.3 ± 1.3 |
|  | H | **5** | 18.9<sub>33</sub> | 27.2 ± 1.8 | 8.8<sub>33</sub> | 21.4<sub>33</sub> |
|  |  | **5a** | 7.2 ± 0.7 | 1.8 ± 0.4 | 6.2 ± 2.4 | **44.2 ± 3.4** |
|  | H | **6** | **82.6 ± 2.2** | 0 | 0 | 0 |
|  |  | **6a** | 14.6 ± 4.8 | 2.0 ± 1.9 | 4.0 ± 0.2 | 6.7 ± 3.2 |
|  |  | **6b** | 30.4 ± 1.2 | 1.0 ± 1.0 | 7.2 ± 3.0 | 21.7 ± 1.8 |
|  |  | **6c** | 7.1 ± 1.8 | 1.9 ± 0.3 | 4.3 ± 2.6 | **78.6 ± 1.8** |
|  | H | **7** | **52.9 ± 3.0** | 0 | 11.3 ± 0.9 | 0 |
|  |  | **7a** | 14.0 ± 3.7 | 11.3 ± 3.5 | 9.1 ± 1.8 | 21.3 ± 1.9 |
|  | H | **8** | **97.3 ± 0.4**<sub>32</sub> | 8.2 | 17.6 ± 1.8<sub>32</sub> | 0<sub>32</sub> |
|  |  | **8a** | 18.3 ± 2.6 | 2.9 ± 0.6 | 7.6 ± 2.4 | 7.7 ± 3.9 |
|  | H | **9** | **93.6 ± 0.9**<sub>32</sub> | N.D. | 10.6 ± 0.7<sub>32</sub> | 22.1 ± 0.2<sub>32</sub> |
|  |  | **9a** | 5.6 ± 1.3 | 1.9 ± 1.9 | 13.7 ± 1.6 | 18.6 ± 0.2 |
|  | H | **10** | **78.9 ± 1.4** | **42.7 ± 0.5** | **78.2 ± 1.7** | 17.1 ± 1.8 |
|  |  | **10a** | 3.2 ± 1.1 | 8.7 ± 1.2 | **41.9 ± 0.8** | 14.4 ± 2.0 |
|  |  | **10b** | 0 | 16.2 ± 1.1 | 32.1 ± 0.9 | 12.9 ± 0.3 |
|  |  | **10c** | 2.5 ± 0.4 | 6.1 ± 2.3 | 21.3 ± 4.2 | 36.6 ± 4.5 |
|  | H | **11** | **94.5 ± 0.8**<sub>34</sub> | N.D. | 17.2 ± 5.2<sub>34</sub> | 3.9 ± 1.8<sub>34</sub> |
|  |  | **11a** | 0.8 ± 0.8 | 3.0 ± 1.2 | 14.7 ± 2.2 | 11.5 ± 2.9 |

<sup>a</sup> The data are presented as % inhibition measured by TEVC technique on A/M2 channels expressed in *Xenopus oocytes*. The values obtained were the mean of at least three experiments after 100 μM for 2 min of treatment. N.D. not determined. See text and Supporting Information for details.

Several of the polycyclic amine, specifically compounds **4, 5**, **6**, **8**, **9**, and **11**, had previously been evaluated against the M2 ion channel and its amantadine-resistant variants.^32–34^ Although most showed activity against the WT M2 channel of the influenza A virus, none exhibited inhibition of the adamantane-resistant mutants M2 L26F, M2 V27A, or M2 S31N (Table 1).

For this reason, and considering that other polycyclic scaffolds have been reported to inhibit adamantane-resistant M2 mutants, ^30,31,32^ herein we synthesized and evaluated two additional polycyclic scaffolds (compounds **7** and **10**) as alternative candidates. Notably, amine **10** was identified as a triple blocker of the M2 WT, L26F, and V27A channels (Table 1).

Replacing the adamantane moiety in compounds **1–3** with alternative polycyclic scaffolds led to compound **6c**, a noradamantane analogue of compound **3**, which showed the most potent inhibition of the M2 S31N mutant. While **6a** and **10** lacked antiviral activity in influenza-infected cells, both compounds showed promising effects against HCoV-229E *in vitro.* DMPK studies revealed that compound **10** combines broad M2 mutant, favourable pharmacokinetics, positioning it as a strong adamantane alternative for antiviral development.

## Results and Discussion

### Chemistry

During the past years, our research groups have developed a series of polycyclic amine analogues based on adamantane scaffold, incorporating various structural modifications. This set included oxadamantane derivatives, ring-rearranged scaffolds, and ring-contracted analogues. Several of these compounds, specifically amine **4**, **5**, **6**, **8**, **9** and **11**, were tested against both the M2 WT channel and its adamantane-resistant variants S31N and V27A.^32–34^ Despite their structural diversity, none of the tested compounds showed significant inhibitory activity against the mutant M2 channels (Table 1), underscoring the challenge of overcoming resistance.

In parallel, and inspired by the S31N activity previously observed for aryl- and heteroaryl-substituted adamantane derivatives such as compounds **1–3**, we adopted a scaffold-hopping approach in which the adamantane core was replaced with alternative polycyclic scaffolds. This strategy aimed to preserve the favorable interactions provided by the aromatic or heteroaromatic substituents, while exploring how variations in the size and shape of the polycyclic core influence activity. By introducing modified cage-like structures, we sought to evaluate the impact of scaffold geometry on channel inhibition and to identify the core dimensions best suited for targeting the M2 WT and mutants channel.

To this end, we first synthesized compounds **4** ^35^, **5** ^33^, **7** ^36^, **8** ^37^, **9** ^37^, **10** ^38^, and **11** ^34^ following previously reported procedures, while compound **6** was obtained from commercial sources. These polycyclic amines were subsequently coupled to aryl groups via a methylene linker using the amino group of the adamantane analogues, through a reductive amination reaction. The synthetic routes of compounds are shown in Scheme 1. We employed a previously described protocol ^16^ with a slight modification: the primary amine was reacted with the corresponding aryl aldehyde in the presence of glacial acetic acid and sodium cyanoborohydride in methanol for 2–3 hours. If unreacted imine was detected, the reaction mixture was stirred for an additional 16 hours. This method afforded *N-*5-(2-bromothiophenyl)methyl derivatives **4b**, **6b**, **9a**, **10b**, and **11a**, as well as *N-*5-[(2-thiophenyl)isoxazolyl]methyl analogues **6c** and **10c**.

For the synthesis of *N*-(2-hydroxy-4-methoxyphenyl)methyl derivatives **5a–8a** and **10a**, the primary amines were initially reacted with the corresponding aldehyde and sodium cyanoborohydride in methanol for 2–3 hours. Since unreacted imines were frequently observed, these intermediates were first purified by column chromatography, then activated by treatment with *p*-toluenesulfonic acid monohydrate (PTSA) ^29^ to generate their protonated forms, and finally reduced with sodium borohydride, yielding the desired secondary amines.

**Scheme 1.**
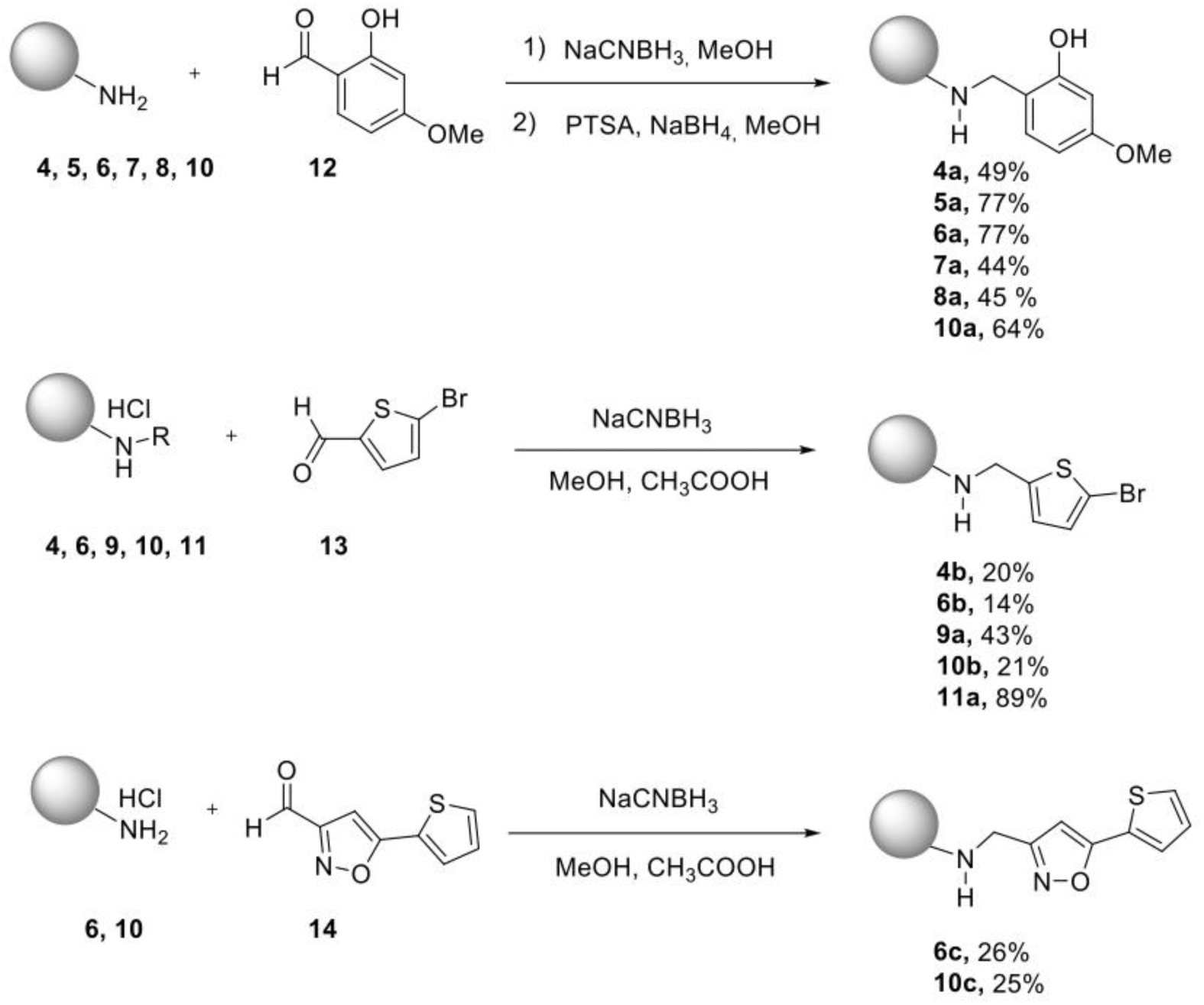
Synthesis of the novel polycyclic derivatives (see text and Supporting Information for details).

### Biology

#### Electrophysiology

The electrophysiological evaluation of the synthesized polycyclic primary amine not tested before (**7** and **10**) and their aryl/heteroaryl derivatives (**4a**-**11a**, **4b**, **6b**, **10b**, **6c** and **10c**) using the two-electrode voltage clamp (TEVC) technique revealed diverse inhibitory profiles against WT and adamantane-resistant A/M2 proton channels (L26F, V27A, and S31N). The inhibitory effects were assessed after 2 minutes of exposure at 100 µM concentration, and the results are summarized in Table 1.^18^ For comparison, we also included in Table 1 the blocking potency of our previously tested amines **4**-**6**, **8**, **9**, **11**^32–34^.

The influence of polycyclic scaffold size and shape on the inhibitory activity against M2 is summarized in Table 1, which clearly demonstrates that subtle variations in ring size, substitution pattern, and conformational flexibility result in distinct activity profiles toward both WT and mutant M2 channels.

Incorporation of a heteroatom into the adamantane ring yield the 2-oxaadamantane (compound **4**), which exhibited strong inhibition of the M2 WT channel (88.2%). In contrast, its regioisomer, **5**, was markedly less effective (18.9%). These results suggest that placing a polar heteroatom on the side of the cage opposite the amine group significantly reduces potency against the WT channel.

The ring-contracted analog 3-noradamantanamine (compound **6**) also demonstrated potent M2 WT inhibition (82.6%), indicating that smaller cage-like scaffolds can effectively mimic adamantane in terms of binding and activity (Figure 1).

The effect of methyl substitution within the polycyclic framework varied depending on both the position of the methyl group and the overall size of the scaffold. In tricyclic derivatives such as compounds **7, 8,** and **9**, the presence of a methyl group near the amine moiety, as in compound **7**, proved detrimental, leading to only moderate inhibition of the M2 WT channel (52.9%). In contrast, compounds **8** and **9**, where steric hindrance around the amine is reduced, retained high levels of inhibition (97.3% and 93.6%, respectively).

Remarkably, tetramethyltetracyclo[4.4.0.0^3.6^.0^4.8^]dec-1-ylamine (compound **10**), a ring-expanded scaffold bearing four methyl groups, —represented the optimal ring size for broad inhibitory activity. Compound **10** not only retained strong inhibition of M2 WT (78.9 ± 1.4%), but also showed moderate activity against L26F (42.7 ± 0.5%) and strong against V27A (78.2 ± 1.7%), making it the only structure among the series capable of efficiently targeting adamantane-resistant channels. These results indicate that a finely balanced ring size—sufficiently large to occupy the hydrophobic channel pocket but not excessively bulky to hinder mutant accommodation—is essential for dual WT/resistant binding.

On the other hand, over-extended derivatives such as compound **11**, bearing a pentacyclic-scaffold, showed excellent WT inhibition (94.5 ± 0.8%) but complete loss of activity against mutants compared to compound **10**, further supporting the notion that excessive scaffold expansion leads to steric incompatibility with the altered mutant pore architecture.

In case of aryl or heteroaryl susbtitution, a marked reduction in potency was observed compared to the corresponding parent amines. For instance, while the unconjugated 2-oxa-adamantane (compound **4**) exhibited high M2 WT inhibition (88 %), its aryl/heteroaryl derivatives **4a** and **4b** displayed significantly decreased activity (23.2 ± 2.4% and 1.0%, respectively). A similar trend was observed with compound **5** (18.9%), whose aryl conjugate **5a** dropped to 7.2 ± 0.7%.

This loss of activity upon arylation was also evident in the tricyclic scaffolds. Compounds **7**, **8**, and **9** showed potent inhibition of M2 WT (52.9–97.3%), yet their respective aryl/heteroaryl derivatives **7a, 8a**, and **9a** exhibited greatly diminished potency (14.0%, 18.3%, and 5.6%, respectively). These findings suggest that aryl substitution at the amine position may disrupt key interactions with the WT channel, either by introducing steric hindrance that impairs proper orientation within the pore or by altering the electronic environment necessary for optimal binding.

Exceptions to this trend were compounds **5a** and **10a**, which showed moderate activity against M2 S31N (44.2 ± 3.4%) and M2 V27A (41.9 ± 0.8%), respectively. Furthermore, notable activity was found with compound **6c**, a 3-noradamantanamine– based heteroaryl derivative, which retained significant inhibitory activity against M2 S31N (78.6 ± 1.8%) while remaining inactive against the WT and other mutant channels. This selective profile suggests that the aryl group in **6c** may exploit a unique pocket or conformational feature specific to the S31N mutant. Notably, **6c** was the most potent S31N-selective inhibitor in the entire series, demonstrating the potential of rational heteroaryl substitution to overcome specific resistance mechanisms.

Altogether, these data indicate that aryl/heteroaryl substitution does not universally enhance M2 channel inhibition and, in most cases, diminishes activity relative to the parent amines.

#### Virus inhibition assay results

The antiviral activity of the most promising compounds (**4, 6, 6c, 7, 8, 9, 10** and **11**) was further evaluated in Madin-Darby canine kidney cells (MDCK) cells infected with two influenza A virus strains: A/PR/8/34 (H1N1), bearing a mutated M2 channel (S31N), and A/HK/7/87 (H3N2), which retains the WT M2 sequence. Both cytopathic effect (CPE) and metabolic assays (MTS) were employed to determine EC₅₀ values after 72 hours of treatment, with corresponding cytotoxicity assessed in uninfected cells (see Table S1, Supporting Information).^16^

Although several compounds demonstrated potent inhibition of the WT M2 channel in the TEVC assay, particularly compounds **4, 6, 8, 9, 10**, and **11**, all of which exhibited >78% inhibition, the majority failed to show antiviral efficacy in infected cells. Notable exceptions include compound **6**, which showed antiviral activity against H3N2 with EC₅₀ values of 26 µM in both CPE and MTS assays. The other exception is compound **8**, one of the most potent M2 WT inhibitors (97.3% inhibition), exhibited strong antiviral activity in H3N2-infected cells, with EC₅₀ values of 6.5 µM (CPE) and 5.9 µM (MTS). Interestingly, compound **7** showed no antiviral effect against the H3N2 strain, despite displaying moderate inhibition of the WT M2 channel in the TEVC assay (52.9%). However, it exhibited the highest potency against the S31N mutant virus (A/PR/8/34), with EC₅₀ values of 4.6 µM (CPE) and 4.4 µM (MTS). Notably, this occurred even though compound **7** did not show measurable activity against the S31N channel in electrophysiological assays, suggesting that its antiviral effect may arise from an alternative mechanism of action unrelated to M2 channel blockade.

Of particular interest, compound **10**, the only structure in our series that effectively blocked both WT and mutant M2 variants (L26F and V27A), failed to demonstrate any antiviral effect. This observation is consistent with previous reports in which M2 channel blockade observed in EP assays did not correlate with antiviral efficacy. It has been proposed that EP recordings taken at early time points (e.g., 2 minutes) are predictive of antiviral activity only for compounds with fast association (fast k_on_) and slow dissociation (slow k_off_) kinetics. In contrast, compounds displaying strong inhibition but no antiviral effect may possess slow binding kinetics (slow k_on_, slow k_off_), making their activity dependent on the equilibrium dissociation constant (K_d_ = k_off_/ k_on_) rather than on the instantaneous percentage of inhibition at short time intervals. ^29, 40^ Altogether, these findings reinforce that M2 inhibition is not a sufficient predictor of antiviral efficacy. Our results align with prior reports and highlight the need to consider both target-based and phenotypic assays when developing antivirals against influenza A.

#### Cytotoxicity evaluation

The cytotoxicity of the most promising compounds was evaluated after 72 hours of treatment in uninfected MDCK cells. Most compounds did not show cytotoxicity with a minimum cytotoxic concentration (MCC) higher than 100 µM. The only exceptions were compounds **6c** and **11**, which exhibited minimum cytotoxic concentrations (MCC) of 0.32 µM and 4 µM, respectively (Table S1, supporting information). Given that compound **6c** is a noradamantane analog of compound **3**, we also evaluated the cytotoxicity of compound **3**, which, despite a previously reported CC₅₀ value of 123 µM in the literature, ^18^ displayed a comparable cytotoxic profile to **6c** in our assay, with an MCC of 0.4 µM. To further investigate whether the cytotoxicity was linked to the 5-(2-thiophenyl)isoxazolyl group present in both **3** and **6c**, we also evaluated the cytotoxicity of compound **10c**, which exhibited moderate cytotoxicity, with an MCC of 8 µM. These cytotoxic effects may partly explain the lack of antiviral activity observed for these compounds *in vitro*. Furthermore, these results suggest that the 5-(2-thiophenyl)isoxazolyl moiety, although previously associated with inhibition of the S31N mutant M2 channel, contribute to undesirable cytotoxic effects. Therefore, while aryl–adamantane substitution has been explored as a strategy for targeting resistant M2 variants, our findings highlight the importance of balancing antiviral potency with cellular safety when optimizing such scaffolds.

#### Activity against HCoV-229E

Amantadine and rimantadine have showed activity against other respiratory viruses, including coronavirus. ^41, 42^ Accordingly, the most promising compounds, **6c** and **10**, were tested for antiviral activity against human coronavirus HCoV-229E using HEL cells.

Both compounds demonstrated activity against HCoV-229E. Notably, compound **10** exhibited a CPE EC₅₀ of 4.7 µM, comparable in potency to the reference compound **GS-441524** (EC₅₀ = 3.8 µM) (see Table S2, Supporting Information). Compound **6c** shown a CPE EC₅₀ of 11 µM. Furthermore, both compounds did not exhibit cytotoxicity in mock-infected cells, as no MCC was detected at concentrations up to 100 µM.

### Drug Metabolism and Pharmacokinetics (DMPK) Assays

Given that compounds **6c** and **10** did not exhibit antiviral activity against influenza virus but demonstrated significant activity against HCoV-229E, further DMPK profiling was undertaken.

Compound **6c** displayed very low metabolic stability in human liver microsomes, with only 0.1% of the parent compound remaining after 60 minutes of incubation (Table 2). In contrast, compound **10** exhibited high metabolic stability, with 94.5% remaining under the same conditions. Based on these results, compound **10** was selected for additional *in vitro* evaluations.

**Table 2.** Microsomal Stability Values of **6c** and **10**.

| Cpd | Microsomal stability |
| --- | --- |
|  | % remanent in human<sup>a</sup> |
| **6c** | 0.1 |
| **10** | 94.5 |

<sup>a</sup>Percentage of remaining compound after 60 min of incubation with pooled human microsomes in the presence of NADPH at 37 °C.

In further *in vitro* profiling, compound **10** demonstrated favorable DMPK characteristics, including high permeability in Caco-2 cell assays, and no inhibition of the hERG potassium channel. Moreover, compound **10** showed no significant inhibition of major cytochrome P450 isoforms, specifically CYP1A2, CYP2C19, and CYP3A4 (Table 3).

**Table 3.**
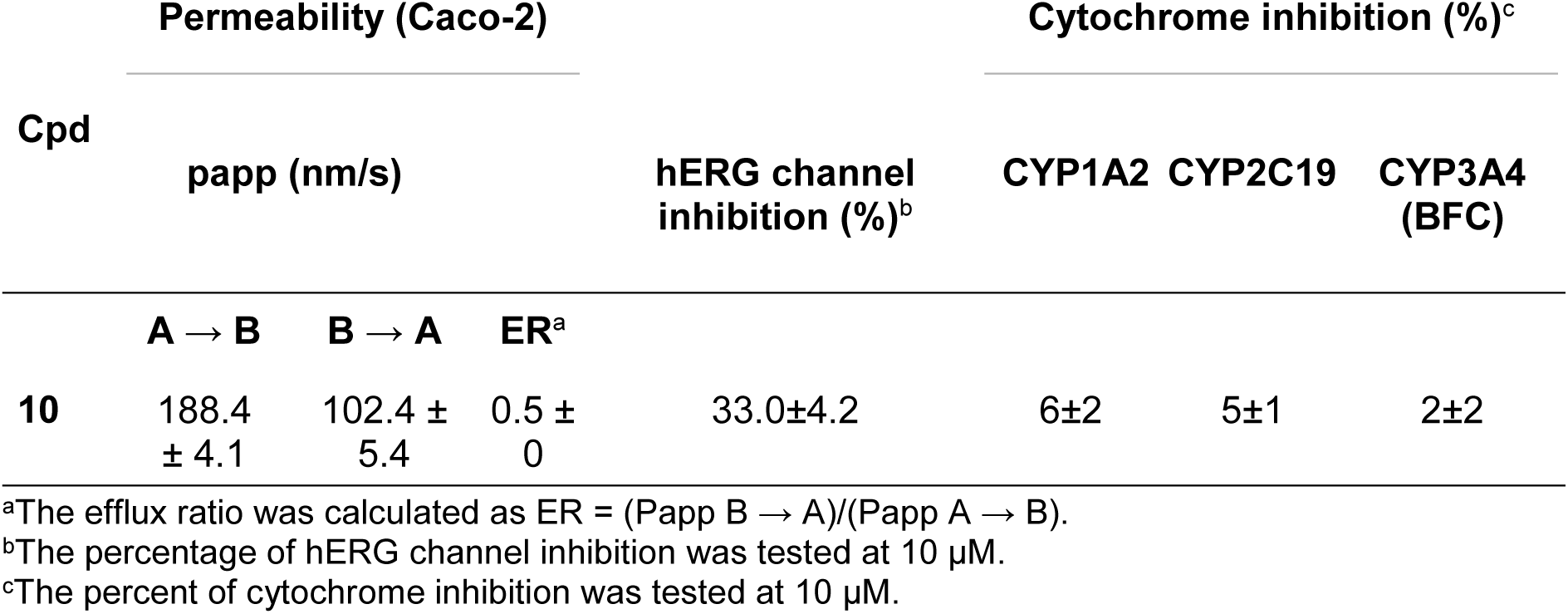
Permeability (PAMPA–BBB) Values, hERG channel inhibition, inhibition of Pooled Human Cytochromes P450 Enzymes of compound 10.

The combination of high permeability and antiviral activity against HCoV-229E suggests that the lack of effect against influenza virus is unlikely to be attributed to permeability or metabolic liabilities. For detailed DMPK assay protocols refer methodology section.

### MD simulations

To gain deeper insight into the role of the polycyclic scaffold in modulating M2 channel inhibition, specifically to rationalize why compound **6c** exhibited inhibitory activity against the S31N mutant whereas compound **10c**, which also bears the 5-(2-thiophenyl)-isoxazolyl moiety, did not, we examined their respective binding modes within the M2(S31N) channel. Representative docking poses were generated for both compounds to evaluate how scaffold-specific differences influence channel interactions and, ultimately, biological activity.

Docking simulations were conducted using the M2(22–46) S31N tetramer. In the complexes formed with M2WJ352 or its analogue **6c** the ammonium group of the ligand was consistently oriented toward the *N*-terminal (extracellular) end of the channel, while the adamantyl moiety was positioned toward the *C*-terminal (intracellular) region. This orientation is in agreement with previously reported data from solution-state NMR in micelles, ^17^ solid-state NMR (ssNMR),^39^ and prior molecular dynamics simulations,^17, 39, 43^ supporting a canonical adamantane-like binding mode within the channel pore of the S31N mutant.

To evaluate the dynamic behavior of ligand–channel complexes, we performed 500 ns MD simulations of the M2(22–46) S31N tetramer embedded in POPC lipid bilayers, in the presence and absence of ligands, using the CHARMM36m force field (see Figure S2, supporting information) ^44, 45^. Representative structures and RMSD plots are shown in Figure 3, while Figure 4 presents three-dimensional water density distributions over the course of the simulation.

**Figure 3.**
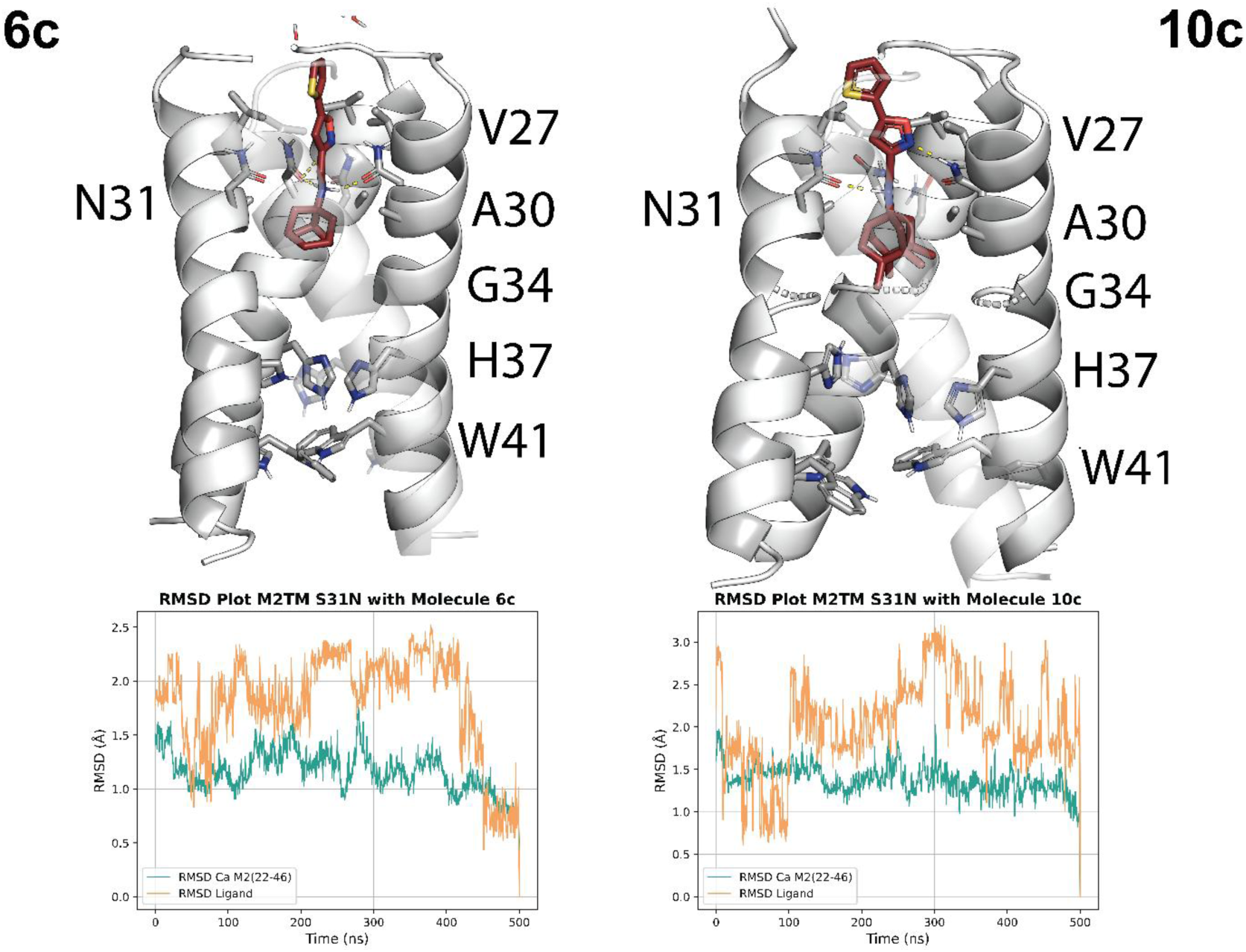
Representative structures and RMSD plots from 500ns-MD simulations of M2(22-46) S31N – drug complexes embedded in POPC bilayers using the CHARMM36m force field. ^44, 45^ (A) **6c**; (B) **10c**. The M2 channel is presented with the cartoon representation in white color, each ligand is colored deep red in stick representation as well as the protein residues in grey color. Water molecules are shown in stick representation with the oxygen atom in red and hydrogen atoms displayed in white color.

**Figure 4.**
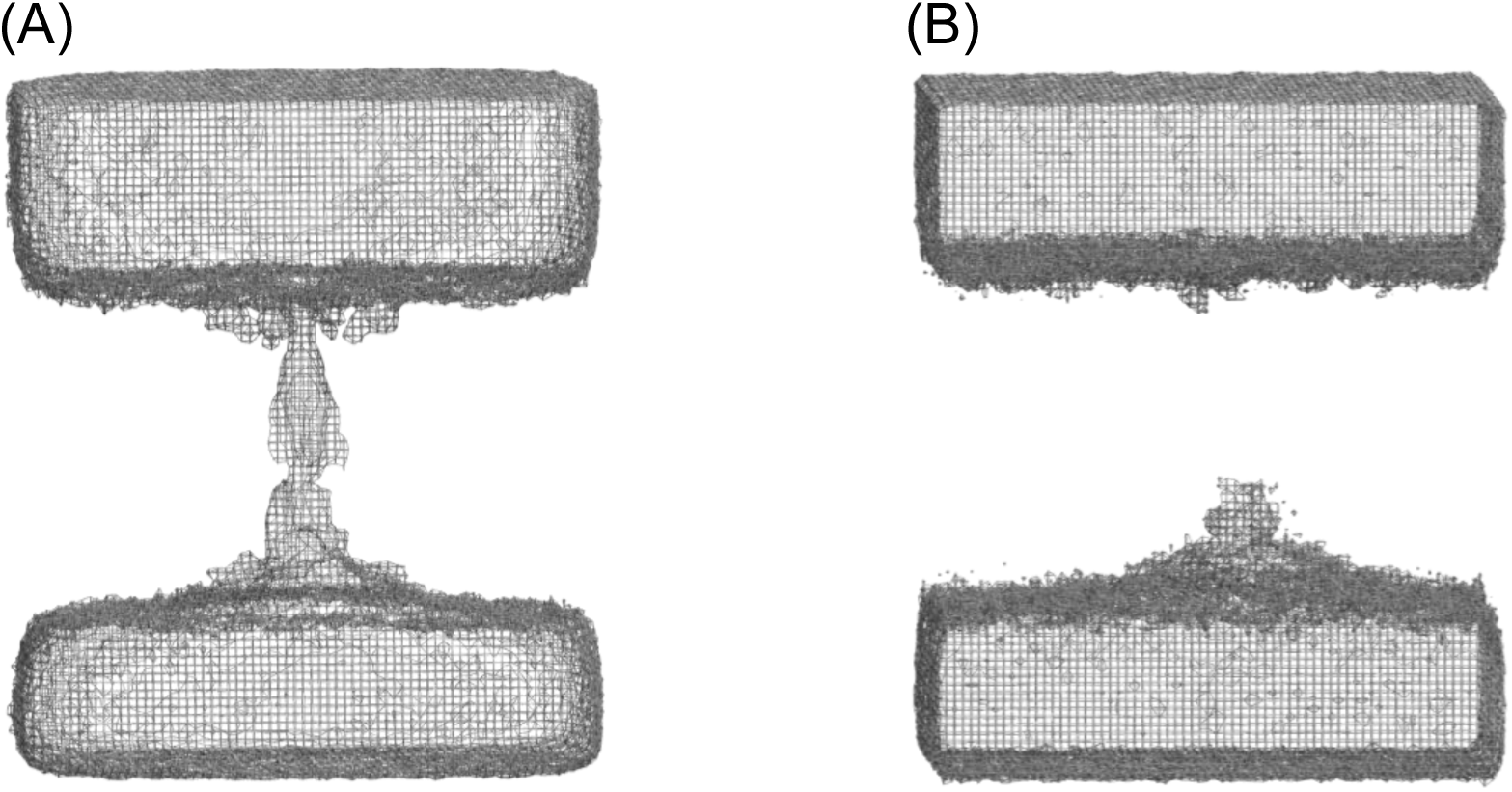
Image of three-dimensional density data of water molecules from the 500 ns MD simulations of M2(22-46) S31N (A) without and (B) with **6c** or **10c** embedded in POPC bilayers using the CHARMM36m force field.^44, 45^

In the apo form, the M2(22–46) S31N channel allowed unrestricted water flux from the Val27 secondary gate to Trp41. By contrast, in the presence of **6c** no water molecules traversed the Val27 gate (Figure 3A and 4A). In this complex, the ammonium group of the ligand forms stable hydrogen bonds with the Asn31 side chains and surrounding water molecules, while the isoxazole ring projects toward the extracellular side, intercalating between the Val27 residues and effectively blocking the water pathway.

Surprisingly, the simulation of the **10c**–M2(S31N) complex also revealed a binding pose consistent with channel occlusion (Figures 3B and 4B), despite the absence of functional inhibition in EP assays. These findings suggest that **10c**, due to its bulkier polycyclic scaffold, may be unable to enter the pore from the *N*-terminal end, as previously supported by ssNMR and liposomal proton flux experiments. The data collectively point to a model in which **10c** acts as a slowly dissociating ligand, resulting in inefficient functional blockade under EP conditions, despite apparent structural engagement with the channel pore.^18, 27,45^

## Conclusions

In this study, we have systematically explored the impact of seven polycyclic cage amines as replacements for the adamantane core in M2 ion channel inhibitors. Our results demonstrate that scaffold size, shape, and substitution influence the ability of these molecules to block WT and resistant forms of the M2 channel. The amine **10** emerged as a triple inhibitor active against M2 WT and the L26F and V27A mutants, while the heteroaryl conjugate **6c** was identified as the most potent S31N-selective inhibitor among the tested series.

However, the disconnect between M2 channel inhibition and antiviral efficacy in cell-based assays underscores the importance of considering binding kinetics during lead optimization. Notably, the observed cytotoxicity of compound **6c** and related isoxazole conjugates, such as the previously reported compound **3**, with MCC below 0.5 µM, may contribute to their limited efficacy and underscores the need to balance antiviral potency with cellular safety.

In contrast, the significant activity of compound **10** against HCoV-229E, together with its favorable DMPK profile, positions it as a promising scaffold for further development. Overall, our findings highlight the importance of scaffold modulation in overcoming drug resistance and extend the potential utility of M2-targeting compounds beyond influenza toward broader antiviral applications.

## Methods

### Chemistry general methods

400 MHz ^1^H NMR / 100.6 MHz ^13^C NMR spectra were recorded on a Varian Mercury 400. The chemical shifts are reported in ppm (δ scale) relative to internal tetramethylsilane, or to solvent peak, and coupling constants are reported in Hertz (Hz). Assignments given for the NMR spectra of the new compounds have been carried out based on heterocorrelation 1H/13C (HSQC) experiments. The used abbreviations were: s, singlet; d, doublet; t, triplet; q, quadruplet; m, multiplet; cs, complex signal; broad s., broad singlet or combinations thereof. All NMR and HPLC-UV spectra scan be found in the Supporting Information. Column chromatography was performed on silica gel 60 Å (Sigma Aldrich, 40 - 63 μm, 230-400 mesh) or with a CombiFlash Rf 150 Teledine ISCO provided with a UV-vis detector. Thin Layer Chromatography (TLC) was performed with aluminium-backed sheets with silica gel 60 F254 (Merck, ref 1.05554), and spots were visualized with UV light. Accurate mass spectra were recorded with ESI techniques on a Hewlett-Packard 5988a LC/MSD-TOF instrument at Unitat d’Espectrometria de Masses dels Centres Científics i Tecnològics de la Universitat de Barcelona (CCiTUB). The analytical samples of all the new compounds, which were subjected to pharmacological evaluation, possessed purity ≥ 95% as evidenced by their HPLC-UV. HPLC-UV were determined with a HPLC Agilent 1260 Infinity II LC/MSD coupled to a photodiode array and mass spectrometer. Samples (5 μL, 0.5 mg/mL) in methanol were injected using an Agilent Poroshell 120 EC-C18 (2.7 μm, 50 mm × 4.6 mm) column at 40 °C. The mobile phase was a mixture of water with 0.05% formic acid (A) and acetonitrile with 0.05% formic acid (B) with a flow 0.6 mL/min, using the following gradients: from 95% A–5% B to 100% B in 3 min; 100% B for 3 min; from 100% B to 95% A–5% B in 1 min; and 95% A–5% B for 3 min. Purity is given as % of absorbance at 254 nm or 275 nm.

### General procedure A

Polycyclic amine (1.2 eq) and aldehyde (1 eq) were mixed in methanol and then treated with NaCNBH_3_ (3 eq). The mixture was stirred at room temperature under argon overnight. The reaction mixture was quenched with H_2_O, and the product was extracted with EtOAc. The combined organic layer was dried over Na_2_SO_4_, filtered and concentrated under reduced pressure. The crude mixture was separated by flash column chromatography (the purification procedure is specified for each compound) to give the target imine. The imine (1 eq) was dissolved in MeOH and to the solution was added *p*-toluenesulfonic acid monohydrate (1 eq) and NaBH_4_ (4 eq) and the mixture was stirred at r.t. for 4 h. The reaction mixture was quenched with saturated aqueous solution of NaHCO_3_ (10 mL) and extracted with DCM. The combined organic extract was dried over anhydrous Na_2_SO_4_, filtered and concentrated under reduced pressure. The mixture was then purified by silica gel flash column chromatography (the gradient used was specified in each compound) to yield the final product.

### General procedure B

To a solution of the amine hydrochloride (1 eq) in methanol (1.5 mL) was added the corresponding aldehyde (1.1 eq), NaCNBH_3_ (2 eq) and glacial acetic acid (2 eq). The reaction mixture was stirred for 3 h at room temperature. Then, two more equivalents of NaCNBH_3_ were added and the reaction was stirred at room temperature overnight. The solvent was removed under vacuum and the resulting residue was partitioned between DCM (5 mL) and water (5 mL). The mixture was basified with 5 N NaOH aqueous solution to basic pH. The layers were separated, and the aqueous layer extracted with further DCM (3 x 5 mL). The combined organics were dried over anhydrous Na_2_SO_4_, filtered and concentrated under reduced pressure to obtain a mixture of the desired product and the alcohol. Column chromatography in silica gel using as eluent hexane to ethyl acetate/hexane mixture gave the amine. Its hydrochloride was obtained by adding an excess of HCl / dioxane to a solution of the amine in EtOAc followed by a filtration under vacuum.

### 2-(((2-Oxaadamantan-1-yl)amino)methyl)-5-methoxyphenol (4a)

To a solution of amine **4** (29 mg, 0.19 mmol) and 2-hydroxy-4-methoxybenzaldehyde (25 mg, 0.16 mmol) in MeOH (2 mL), NaCNBH3 (23 mg, 0.36 mmol) was added following the general procedure A. Purification by Combiflash® in silica gel using as eluent hexane to ethyl acetate/hexane mixture (8/2) gave the imine as a yellow oil (21 mg, 46 % yield). Reaction of the imine (21 mg, 0.07 mmol), PTSA (13 mg, 0.07 mmol), NaBH4 (11 mg, 0.28 mmol) in MeOH (2 mL) afforded the title compound as a white solid (10 mg, 49 % yield). 1H NMR (400 MHz, CDCl3) δ: 1.60 [dm, J = 12.5 Hz, 2H, 4(10)-Ha], 1.77-1.82 [cs, 4H, 6-H2 and 8(9)-Ha], 1.87-1.94 [cs, 4H, 4(10)-Hb and 8(9)-Hb], 2.25 [s, 2H, 5(7)-Hb], 3.74 (s, 3H, OCH3), 4.08 (s, 2H, NH-CH2), 4.27 (br s, 1H, 3-H), 6.38 (dd, 1H, J = 8.3 Hz, J’ = 2.4 Hz, 4’-H), 6.54 (s, 1H, 6’-H), 7.01 (d, J = 8.3 Hz, 3’-H).

13C-NMR (100.6 MHz, CDCl3) δ: 28.0 [CH, C5(7)], 34.68 (CH2, C6), 34.71 [CH2, C4(10)], 38.8 [CH2, C8(9)], 41.9 (CH2, NH-CH2), 55.4 (CH3, OCH3), 71.4 (CH, C3), 83.4 (C, C1), 103.1 (CH, C6’), 106.4 (CH, C4’), 114.6 (C, C2’), 130.6 (CH, C3’), 158.0 (C, C1’), 161.0 (C, C5’). HRMS-ESI+ m/z [M+H]+ calcd for [C17H24NO3]+: 290.1751, found: 290.1759. Purity by HPLC at 275 nm: 95.04%.

### *N*-((5-Bromothiophen-2-yl)methyl)-2-oxaadamantan-1-amine hydrochloride (4b)

Following the general procedure B, 2-oxaamantadine hydrochloride (200 mg, 1.05 mmol), 5-bromothiophene-2-carbaldehyde (130 mg, 1.16 mmol), NaCNBH_3_ (264 mg, 4.2 mmol) and glacial acetic acid (0.12 mL, 2.10 mmol) were mixed to obtain a yellowish oil (468 mg). After column chromatography in silica gel **4b** was obtained as a white oil (68 mg, 20 % yield) that formed its hydrochloride salt as a white solid (81 mg). ^1^H NMR (400 MHz, MeOD) *δ*: 1.76 [dm, *J* = 12.4 Hz, 2H, 4(10)-Ha], 1.89 (dm, *J* = 12.8 Hz, 1H, 6-Ha), 1.95-2.04 [cs, 5H, 6-Hb and 8(9)-H_2_], 2.12 (d, *J* = 11.2 Hz, 2H, 4(10)-Hb], 2.40 [s, 2H, 5(7)-H], 4.37 (broad s, 1H, 3-H), 4.46 (s, 2H, NH-C<u>H_2_</u>), 7.09-7.11 (cs, 2H, 3’-H and 4’-H). ^13^C-NMR (100.6 MHz, MeOD) δ: 29.3 [CH, C5(7)], 34.93 (CH_2_, C6), 34.98 [CH_2_, C4(10)], 37.8 [CH_2_, C8(9)], 39.0 (CH_2_, <u>C</u>H_2_-NH), 73.7 (CH, C3), 86.7 (C, C1), 115.6 (C, C2’), 131.8 (CH, C3’), 132.5 (CH, C4’), 135.9 (C, C5’). HRMS-ESI+ *m/z* [M+H]^+^ calcd for [C_14_H_19_BrNOS]^+^: 328.0365, found: 328.0363. Purity by HPLC at 254 nm: 97.88%.

### 2-(((2-Oxaadamantan-5-yl)amino)methyl)-5-methoxyphenol (5a)

To a solution of amine **5** (22 mg, 0.14 mmol) and 2-hydroxy-4-methoxybenzaldehyde (19 mg, 0.12 mmol) in MeOH (2 mL), NaCNBH_3_ (23 mg, 0.36 mmol) was added following the general procedure A. Purification by Combiflash® in silica gel using as eluent hexane to ethyl acetate/hexane mixture (8/2) gave the imine as a yellow solid (22 mg, 65 % yield). Reaction of the imine (22 mg, 0.08 mmol), PTSA (15 mg, 0.08 mmol), NaBH_4_ (12 mg, 0.32 mmol) in MeOH (2 mL) afforded the title compound as a white solid (17 mg, 77 % yield). ^1^H NMR (400 MHz, CDCl_3_) *δ*: 1.54 [dm, *J* = 12.4 Hz, 2H, 8(10)-Ha], 1.67 [d, *J* = 12 Hz, 2H, 4(9)-Ha), 1.86 [broad s, 2H, 6-H_2_], 1.96-2.02 [cs, 2H, 4(9)-Hb and 8(10)-Hb], 2.29 (s, 1H, 7-H), 3.76 (s, 3H, OC<u>H_3_</u>], 3.93 (s, 2H, NH-C<u>H_2_</u>), 4.22 [br s, 1H, 1(3)-H), 6.34 (dd, 1H, *J* = 8.2 Hz, *J’* = 2.4 Hz, 4’-H), 6.41 (d, *J* = 2.4 Hz, 1H, 6’-H), 6.87 (d, *J* = 8.2 Hz, 3’-H). ^13^C-NMR (100.6 MHz, CDCl_3_) δ: ^13^C-NMR (100.6 MHz, CDCl_3_) δ: 27.4 (CH, C7), 35.1 [CH_2_, C8(10)], 40.6 (CH_2_, C6), 41.0 [CH_2_, C4(9)], 43.4 (CH_2_, NH-<u>C</u>H_2_), 50.6 (C, C5), 55.4 (CH_3_, OCH_3_), 69.1 [CH, C1(3)], 102.3 (CH, C6’), 105.1 (CH, C4’), 115.7 (C, C2’), 128.6 (CH, C3’), 159.4 (C, C1’), 160.5 (C, C5’). HRMS-ESI+ *m/z* [M+H]^+^ calcd for [C_17_H_24_NO_3_]^+^: 290.1751, found: 290.1758.

### 2-(((Tricyclo[3.3.1.0^3,7^]non-3-yl)amino)methyl)-5-methoxyphenol hydrochloride (6a)

To a solution of amine **6** (70 mg, 0.51 mmol) and 2-hydroxy-4-methoxybenzaldehyde (64 mg, 0.43 mmol) in MeOH (6 mL) NaCNBH_3_ (80 mg, 1.28 mmol) was added following the general procedure A. Purification by Combiflash® in silica gel using as eluent hexane to ethyl acetate/hexane mixture (4/6) gave the imine as a yellow solid (99 mg, 86% yield). The imine (99 mg, 0.36 mmol), PTSA (70 mg, 0.36 mmol) and NaBH_4_ (55 mg, 1.46 mmol) in MeOH (2 mL) afforded a crude that after purification by Combiflash® in silica gel using as eluent dichloromethane to methanol/dichloromethane mixture (1/9) yielded the desire product as a white solid (76 mg, 77 % yield). ^1^H NMR (400 MHz, CDCl_3_) *δ*: 1.53 (m, 2H, 9-Ha), 1.58 (broad d, *J* = 11.2 Hz, 2H, 6(8)-Ha), 1.64 (m, 1H, 9-Hb), 1.83 [cs, 4H, 2(4)-H_2_], 1.85-1.89 (m, 2H, 6(8)-Hb), 2.21 (tt, *J* = 6.8 Hz, *J’* = 1.8 Hz, 1H, 7-H), 2.30 (br s, 2H, 1(5)-H), 3.76 (s, 3H, OC<u>H_3_</u>), 3.94 (s, 2H, NH-C<u>H_2_</u>), 6.33 (dd, *J* = 8.2 Hz, *J’* = 2.6 Hz, 1H, 4’-H), 6.40 (d, *J* = 2.6 Hz, 1H, 6’-H), 6.87 (d, *J* = 8.2 Hz, 1H, 3’-H). ^13^C-NMR (100.6 MHz, CDCl_3_) δ: 34.9 (CH_2_, C9), 37.6 [CH, C1(5)], 42.6 (CH, C7), 43.7 [CH_2_, C6(8)], 47.3 (CH_2_, <u>C</u>H_2_-NH), 48.4 [CH_2_, C2(4)], 55.4 (CH_3_, OCH_3_), 68.5 (C, C3), 102.3 (CH, C6’), 104.9 (CH, C4’), 116.0 (C, C2’), 128.6 (CH, C3’), 159.7 (C, C1’), 160.4 (C, C5’). HRMS-ESI+ *m/z* [M+H]^+^ calcd for [C_17_H_24_NO_2_]^+^: 274.1802, found: 274.1808. Purity by HPLC at 275 nm: 98.98%.

### *N*-((5-Bromothiophen-2-yl)methyl)tricyclo[3.3.0.0^3,7^]nonan-1-amine hydrochloride (6b)

Following the general procedure B, 3-noradamantanamine hydrochloride (120 mg, 0.69 mmol), 5-bromothiophene-2-carbaldehyde (145 mg, 0.76 mmol), NaCNBH_3_ (173 mg, 2.76 mmol) and glacial acetic acid (0.08 mL, 1.38 mmol) were mixed to obtain a yellow oil (313 mg). After column chromatography a white solid was obtained (30 mg, 14 % yield) that formed its hydrochloride salt as a white solid (35 mg). ^1^H NMR (400 MHz, MeOD) *δ*: 1.61 (dt, *J* = 13.3 Hz, *J’* = 2.7 Hz, 1H, 9-Ha), 1.70-1.76 [cs, 3H, 6(8)-Ha and 9-Hb], 1.99-2.01 [m, 2H, 6(8)-Hb], 2.05-2.11 [cs, 4H, 2(4)-H_2_], 2.43 [broad s, 2H, 1(5)-H], 2.46 (m, 1 H, 7-H), 4.40 (s, 2H, NH-C<u>H_2_</u>), 7.12-7.14 (cs, 2H, 3’-H and 4’-H). ^13^C-NMR (100.6 MHz, MeOD) δ: 34.9 (CH_2_, C9), 38.8 [CH, C1(5)], 42.6 (CH_2_, <u>C</u>H_2_-NH), 43.67 (CH, C7), 43.70 [CH_2_, C6(8)], 46.3 [CH_2_, C2(4)], 71.8 (C, C3), 115.6 (C, C2’), 131.9 (C, C3’ or C4’), 132.4 (C, C4’ or C3’), 136.1 (C, C5’). HRMS-ESI+ *m/z* [M+H]^+^ calcd for [C_14_H_19_BrNS]^+^: 312.0416, found: 312.0421. Purity by HPLC at 254 nm: 99.50%.

### *N*-((5-(Thiophen-2-yl)isoxazol-3-yl)methyl)tricyclo[3.3.0.0^3,7^]nonan-1-amine hydrochloride (6c)

A solution of **6**·HCl (50 mg, 0.29 mmol) in MeOH (6 mL) was prepared in a round bottom flask equipped with a CaCl_2_ tube. To that, NaBH_3_CN (38 mg, 0.35 mmol), acetic acid (0.02 mL, 0.58 mmol) and 5-(thiophen-2-yl)isoxazole-3-carbaldehyde (62 mg, 0.35 mmol) were added. The mixture was stirred at room temperature for 2 h. Then, more NaBH_3_CN (18 mg, 0.28 mmol) and 5-(thiophen-2-yl)isoxazole-3-carbaldehyde (40 mg, 0.22 mmol) were added, and the reaction mixture was stirred at room temperature for another 16 h. The mixture was concentrated under reduced pressure and to the obtained crude was added water (20 mL). Then, NaOH 2N solution was added until basic pH was reached, and the aqueous phase was extracted with DCM (3 x 10 mL), dried over anh. Na_2_SO_4_, filtered and concentrated under vacuum. Its hydrochloride was obtained by adding an excess of HCl / diethyl ether to a solution of the compound in ethyl acetate, followed by filtration to obtain the hydrochloride salt as a white solid (25 mg, 26% yield). The analytical sample was obtained by crystallization from methanol/diethyl ether. ^1^H NMR (400 MHz, MeOD) *δ*: 1.63 (dt, *J* = 13.2Hz, *J’* = 2.4 Hz, 1H, 9-Ha), 1.72-1.77 [cs, 3H, 6(8)-Ha and 9-Hb], 2.02-2.13 [cs, 6H, 2(4)-H_2_ and 6(8)-Hb], 2.46 (br s, 2H, 1(5)H), 2.50 (tt, *J* = 6.8 Hz, *J’* = 2 Hz, 1H, 7-H), 4.40 (s, 2H, NH-C<u>H_2_</u>), 6.78 (s, 1H, 4’-H), 7.21 (dd, *J* = 5.2 Hz, *J’* = 4.0 Hz, 1H, 4’’-H), 7.65 (dd, *J* = 3.6 Hz, *J’* = 1.2 Hz, 1H, 3’’-H), 7.69 (dd, *J* = 5.2 Hz, *J’* = 1.2 Hz, 1H, 5’’-H). ^13^C-NMR (100.6 MHz, MeOD) δ: 34.9 (CH_2_, C9), 38.8 [CH, C1(5)], 40.0 (CH_2_, <u>C</u>H_2_-NH), 43.6 (CH, C7), 43.7 [CH_2_, C6(8)], 46.2 [CH_2_, C2(4)], 72.2 (C, C3), 99.7 (CH, C4’), 129.1 (CH, C3’’), 129.4 (C, C2’’), 129.5 (CH, C4’’), 130.4 (CH, C5’’), 158.8 (C, C3’), 167.9 (C, C5’). HRMS-ESI+ *m/z* [M+H]^+^ calcd for [C_17_H_21_N_2_OS]^+^: 301.1369, found: 301.1373. Purity by HPLC at 254 nm: 99.68%.

### 5-Methoxy-2-((((3,7,9,9-tetramethyl)tricyclo[3.3.0.0^3,7^]non-1-yl)amino)methyl)phenol hydrochloride (7a)

To a solution of amine **7** (25 mg, 0.13 mmol) and 2-hydroxy-4-methoxybenzaldehyde (16 mg, 0.11 mmol) in MeOH (2 mL) NaCNBH_3_ (23 mg, 0.36 mmol) was added following the general procedure A. Purification by Combiflash® in silica gel using as eluent hexane to ethyl acetate/hexane mixture (3/7) gave the imine as a yellow solid (23 mg, 70 % yield). The imine (23 mg, 0.07 mmol), PTSA (13 mg, 0.07 mmol), NaBH_4_ (11 mg, 0.28 mmol) in MeOH (2 mL) afforded a crude that after the purification by Combiflash® in silica gel using as eluent dichloromethane to methanol/dichloromethane mixture (1/9) yielded the desire product as a white solid. Its hydrochloride was obtained by adding an excess of HCl / diethyl ether to a solution of the amine in dichloromethane, concentrated under reduced pressure to give the desired compound (10 mg, 44% yield). ^1^H NMR (400 MHz, MeOD) *δ*: 1.10 [s, 6H, 3(7)-C<u>H_3_</u>), 1.26 [s, 6H, 9-(C<u>H_3_</u>)_2_], 1.42 [dt, 2H, *J* = 10.8 Hz, *J’* = 2.8 Hz, 4(6)-Ha], 1.80 (t, 1H, *J* = 3.2 Hz, 5-H), 1.85 [dd, 2H, *J* = 10.4 Hz, *J’* = 2.4, 4(6)-Hb], 1.89-1.92 [complex signal, 2H, 2(8)-Ha], 2.00 [d, 2H, *J* = 12.8 Hz, 2(8)-Hb], 3.77 (s, 3H, OCH_3_), 4.14 (s, 2H, CH_2_NH), 6.48-6.50 [cs, 2H, 4’(6’)-H], 7.19 (d, 1H, *J* = 8.8, 3’-H). ^13^C-NMR (100.6 MHz, MeOD) 21.8 [CH_3_, C3(7)-<u>C</u>H_3_], 23.8 [CH_3_, C9-(<u>C</u>H_3_)_2_], 39.2 [C, C3(7)], 45.08 (C, C9), 45.14 (CH_2_, <u>C</u>H_2_-NH), 46.5 [CH_2_, C4(6)], 47.9 [CH_2_, C2(8)], 49.6 (CH, C5), 55.8 (CH_3_, OCH_3_), 68.9 (C, C1), 102.3 (CH, C6’), 106.5 (CH, C4’), 111.5 (C, C2’), 133.0 (CH, C3’), 158.2 (C, C1’), 163.6 (C, C5’). HRMS-ESI+*m/z* [M+H]^+^ calcd for [C_21_H_32_NO_2_]^+^: 330.2428, found: 330.2434. Purity by HPLC at 275 nm: 98.15%.

### 2-((((3,7-Dimethyltricyclo[3.3.0.0^3,7^]octan-1-yl)methyl)amino)methyl)-5-methoxyphenol hydrochloride (8a)

To a solution of amine **8** (21 mg, 0.13 mmol) and 2-hydroxy-4-methoxybenzaldehyde (18 mg, 0.12 mmol) in MeOH (2 mL) NaCNBH_3_ (22 mg, 0.35 mmol) was added following the general procedure A. Purification by Combiflash® in silica gel using as eluent hexane to ethyl acetate/hexane mixture (1/9) gave the imine as a yellow solid (10 mg, 41 % yield). The imine (10 mg, 0.033 mmol), PTSA (6 mg, 0.03 mmol), and NaBH_4_ (5 mg, 0.13 mmol) in MeOH (2 mL) afforded a crude that after the purification by Combiflash® in silica gel using as eluent dichloromethane to methanol/dichloromethane mixture (1/9) yielded the desire product as a white solid. Its hydrochloride was obtained by adding an excess of HCl / diethyl ether to a solution of the amine in dichloromethane, concentrated under reduced pressure to give the desired compound (5 mg, 45 % yield). ^1^H NMR (400 MHz, MeOD) *δ*: 1.18 [s, 6H, 3(7)-C<u>H_3_</u>), 1.37 [dd, 2H, *J* = 8.2 Hz, *J’* = 3.2 Hz, 4(6)-Ha], 1.41-1.50 [cs, 4H, 2(8)-H_2_], 1.60 [dd, 2H, *J* = 8.2 Hz, *J’* = 2.8 Hz, 4(6)-Hb], 2.15 (t, 1H, *J* = 2.8 Hz, 5-H), 3.13 (s, 2H, NHCH_2_), 3.76 (s, 3H, OCH_3_), 4.12 (s, 2H, CH_2_NH), 6.44-6.46 (complex signal, 2H, 4’-H, 6’-H), 7.15 (d, 1H, *J* = 7.2 Hz, 3’-H). ^13^C-NMR (100.6 MHz, MeOD) 16.8 [CH_3_, C3(7)-<u>C</u>H_3_], 43.4 (CH, C5), 49.7 (CH_2_, <u>C</u>H_2_-NH), 51.1 (CH_2_, NH-CH_2_), 54.6 [CH_2_, C4(6)], 55.7 (CH_3_, OCH_3_), 57.1 [CH_2,_ C2(8)] 102.4 (CH, C6’), 106.1 (CH, C4’), 112.0 (C, C2’), 132.9 (CH, C3’), 159.3 (C, C1’), 163.4 (C, C5’). HRMS-ESI+ *m/z* [M+H]^+^ calcd for [C_19_H_28_NO_2_]^+^: 302.2115, found: 302.2122. Purity by HPLC at 275 nm: 97.20%.

### *N*-((5-Bromothiophen-2-yl)methyl)-3,7-dimethyltricyclo[3.3.0.0^3,7^]octan-1-amine hydro-chloride (9a)

Following the general procedure B, amine **9**·HCl (110 mg, 0.59 mmol), 5-bromothiophene-2-carbaldehyde (122 mg, 0.64 mmol), NaCNBH_3_ (148 mg, 2.36 mmol) and glacial acetic acid (0.07 mL, 1.18 mmol) were mixed to obtain a yellowish oil (251 mg). After column chromatography a white solid was obtained (83 mg, 43% yield) that formed its hydrochloride salt as a white solid (80 mg). ^1^H NMR (400 MHz, MeOD) *δ*: 1.22 [s, 6H, 3(7)-C<u>H_3_</u>], 1.48 (d, 2H, *J* = 8.8 Hz, 4(6)-Ha], 1.78 [dd, 2H, *J* = 8.8 Hz, *J’* = 2.9 Hz, 4(6)-Hb], 1.82 [broad s, 4H, 2(8)-H_2_], 2.51 (t, *J* = 2.9 Hz, 1H, 5-H), 4.40 (s, 2H, NH-C<u>H_2_</u>), 7.11 (d, *J* = 4.0 Hz, 1H, 3’-H), 7.15 (d, *J* = 4.0 Hz, 1H, 4’-H). ^13^C-NMR (100.6 MHz, MeOD) δ: 16.4 [CH_3_, C3(7)-<u>C</u>H_3_], 43.4 (CH_2_, <u>C</u>H_2_-NH), 43.5 (CH, C5), 47.8 [C, C3(7)], 53.6 [CH_2_, C4(6)], 54.8 [CH_2_, C2(8)], 68.0 (C, C1), 115.5 (C, C2’), 131.8 (CH, C3’), 132.5 (CH, C4’), 136.0 (C, C5’). HRMS-ESI+ *m/z* [M+H]^+^ calcd for [C_15_H_21_BrNS]^+^: 326.0573, found: 326.0575. Purity by HPLC at 254 nm: 99.03%.

### 5-Methoxy-2-(((3,4,8,9-tetramethyltetracyclo[4.4.0.0^3,6^.0^4,8^]decan-1-yl)amino)methyl) phenol (10a)

To a solution of amine **10** (35 mg, 0.17 mmol) and 2-hydroxy-4-methoxybenzaldehyde (22 mg, 0.14 mmol) in MeOH (2 mL), NaCNBH_3_ (27 mg, 0.43 mmol) was added following the general procedure A. Purification by Combiflash® in silica gel using as eluent hexane to ethyl acetate/hexane mixture (1/9) gave the imine as a yellow solid (25 mg, 52 % yield). The imine (25 mg, 0.36 mmol), PTSA (14 mg, 0.07 mmol) and NaBH_4_ (11 mg, 0.29 mmol) in MeOH (2 mL) afforded a crude that after the purification by Combiflash® in silica gel using as eluent dichloromethane to methanol/dichloromethane mixture (1/9), yielded the desire product as a white solid (16 mg, 64 % yield). ^1^H NMR (400 MHz, CDCl_3_) *δ*: 0.78-0.83 [cs, 4H, 2(10)-Ha and 5(7)-Ha], 0.937 [s, 6H, 3(9)-CH_3_ or 4(8)-CH_3_], 0.943 [s, 6H, 4(8)-CH_3_ or 3(9)-CH_3_], 1.80-1.83 [cs, 4H, 2(10)-Hb and 5(7)-Hb], 2.33 (s, 1H, 6-H), 3.76 (s, 3H, OC<u>H_3_</u>), 4.0 (s, 2H, NH-C<u>H_2_</u>), 6.33 (dd, *J* = 8.3 Hz, *J’* = 2.6 Hz, 1H, 4’-H), 6.41 (d, *J* = 2.7 Hz, 1H, 6’-H), 6.88 (d, *J* = 8.2 Hz, 1H, 3’-H). ^13^C-NMR (100.6 MHz, CDCl_3_) δ: 15.6 [CH_3_, C3(9)-CH_3_ or C4(8)-CH_3_], 15.9 [CH_3_, C4(8)-CH_3_ or C3(9)-CH_3_], 39.0 [CH_2_, C5(7)], 40.1 (CH, C6), 42.7 [CH_2_, C2(10)], 44.79 [C, C3(9) or C4(8)], 44.86 [C, C4(8) or C3(9)], 46.8 (CH_2_, NH-<u>C</u>H_2_), 55.4 (CH_3_, OCH_3_), 61.1 (C, C1), 102.3 (CH, C6’), 105.0 (CH, C4’), 128.6 (CH, C3’), 160.5 (C, C5’). Quaternary carbon atoms C1’ and C2’ were not visible in the ^13^C NMR spectrum. HRMS-ESI+ *m/z* [M+H]^+^ calcd for [C_22_H_32_NO_2_]^+^: 342.2428, found: 342.2426. Purity by HPLC at 275 nm: 97.34%.

### *N*-((5-bromothiophen-2-yl)methyl)-3,4,8,9-tetramethyltetracyclo[4.4.0.0^3,6^.0^4,8^]decan-1-amine hydrochloride (10b)

To a suspension of **10**·HCl (60 mg, 0.56 mmol) in toluene (5 mL) was added 5-bromothiophene-2-carbaldehyde (107 mg, 0.56 mmol) and *p*-toluenesulfonic acid (21 mg, 0.11 mmol). The mixture was heated to reflux with a Dean-Stark apparatus overnight. The reaction mixture was allowed to cool down to room temperature and concentrated under vacuum to give a mixture of the imine and (5-bromothiophene-2-yl)methanol as a brown oil (115 mg). This crude was used without purification in the next step.

To a solution of the crude from the previous reaction (115 mg) in glacial acetic acid (5 mL) was added NaCNBH_3_ (426 mg, 6.78 mmol) in small portions in an inert atmosphere. The resulting suspension was stirred at room temperature for 4 h. The reaction mixture was cooled to 0 °C with an ice bath, treated with 10 N NaOH aqueous solution to basic pH and extracted with DCM (3 x 10 mL). The combined organics were dried over anhydrous Na_2_SO_4_, filtered and concentrated under reduced pressure to obtain a mixture of the desired product and (5-bromothiophene-2-yl)methanol. Column chromatography (Hexane/Ethyl acetate mixture) gave a white oil (45 mg, 21% yield), that was dissolved in EtOAc. HCl / dioxane was added to the solution to form the corresponding hydrochloride salt that was obtained after filtration (11 mg). ^1^H NMR (400 MHz, MeOD) *δ*: 0.97 [d, *J* = 12.0 Hz, 2H, 5(7)-Ha], 1.02 [s, 6H, 3(9)-CH_3_ or 4(8)-CH_3_], 1.04 [s, 6H, 4(8)-CH_3_ or 3(9)-CH_3_], 1.10 [d, *J* = 10.6 Hz, 2H, 2(10)-Ha], 1.95 [d, *J* = 12.0 Hz, 2H, 5(7)-Hb], 2.07 [d, *J* = 10.6 Hz, 2(10)-Hb], 2.49 (s, 1H, 6-H), 4.48 (s, 2H, NH-C<u>H_2_</u>), 7.12-7.16 (cs, 2H, 3’-H and 4’-H). ^13^C-NMR (100.6 MHz, CDCl_3_) δ: 15.4 [CH_3_, C3(9)-CH_3_ or C4(8)-CH_3_], 15.6 [CH_3_, C4(8)-CH_3_ or C3(9)-CH_3_], 39.0 (CH, C6), 39.1 [CH_2_, C5(7)], 40.4 [CH_2_, C2(10)], 42.0 (CH_2_, NH-<u>C</u>H_2_), 45.9 [C, C3(9) or C4(8)], 46.1 [C, C4(8) or C3(9)], 64.6 (CH C1), 115.6 (C, C2’), 131.9 (CH, C3’ or C4’), 132.4 (CH, C4’ or C3’), 136.2 (C, C5’). HRMS-ESI+ *m/z* [M+H]^+^ calcd for [C_19_H_27_BrNS]^+^: 380.1042, found: 380.1051. Purity by HPLC at 254 nm: 96.75%.

### 3,4,8,9-Tetramethyl-N-((5-(thiophen-2-yl)isoxazol-3-yl) methyl)tetracyclo[4.4.0.0^3,6^.0^4,8^]decan-1-amine hydrochloride (10c)

A solution of **10**·HCl (28 mg, 0.12 mmol) in MeOH (6 mL) was prepared in a round bottom flask equipped with a CaCl_2_ tube. To that, NaBH_3_CN (15 mg, 0.23 mmol), acetic acid (0.01 mL, 0.24 mmol) and 5-(thiophen-2-yl)isoxazole-3-carbaldehyde (25 mg, 0.14 mmol) were added. The mixture was stirred at room temperature for 2 h. Then, more NaBH_3_CN (8 mg, 0.12 mmol) and 5-(thiophen-2-yl)isoxazole-3-carbaldehyde (16 mg, 0.09 mmol) were added, and the reaction mixture was stirred at room temperature for another 16 h. The mixture was concentrated under reduced pressure and to the obtained crude was added water (20 mL). Then, NaOH 2N solution was added until basic pH was reached, and the aqueous phase was extracted with DCM (3 x 10 mL), dried over anh. Na_2_SO_4_, filtered and concentrated under vacuum. The crude was purified by Combiflash® in silica gel using as eluent diethyl ether to methanol/diethyl ether mixture (1/9), yielded the desire product as a colorless oil (12 mg, 25 % yield). Its hydrochloride was obtained by adding an excess of HCl / diethyl ether to a solution of the compound in ethyl acetate, followed by evaporation to obtain the hydrochloride salt as a white solid. ^1^H NMR (400 MHz, MeOD) *δ*: 0.98 [dd, *J* = 12.0 Hz, *J’* = 3.2 Hz, 5(7)-Ha], 1.03 [s, 6H, 3(9)-CH_3_ or 4(8)-CH_3_], 1.06 [s, 6H, 4(8)-CH_3_ or 3(9)-CH_3_], 1.14 [d, *J* = 10.8 Hz, 2H, 2(10)-Ha], 1.98 [dd, *J* = 12 Hz, *J’* = 1.6 Hz, 2H, 5(7)-Hb], 2.10 [d, *J* = 10.8 Hz, 2H, 2(10)-Hb], 2.50 (broad s., 1H, 6-H), 4.47 (s, 2H, NH-C<u>H_2_</u>), 6.78 (s, 1H, 4’-H), 7.21 (dd, *J* = 5.1 Hz, *J’* = 3.7 Hz, 1H, 4’’-H), 7.65 (dd, *J* = 3.7 Hz, *J’* = 1.2 Hz, 1H, 3’’-H), 7.69 (dd, *J* = 5.2 Hz, *J’* = 1.2 Hz, 1H, 5’’-H). ^13^C-NMR (100.6 MHz, MeOD) δ: 15.4 [CH_3_, C3(9)-CH_3_ or C4(8)-CH_3_], 15.7 [CH_3_, C4(8)-CH_3_ or C3(9)-CH_3_], 39.0 (CH, C6), 39.1 [CH_2_, C5(7)], 39.4 (CH_2_, <u>C</u>H_2_-NH), 40.4 [CH_2_, C2(10)], 45.9 [C, C3(9) or C4(8)], 46.2 [C, C4(8) or C3(9)], 64.8 (CH C1), 99.7 (CH, C4’), 129.1 (CH, C3’’), 129.47 (C, C2’’), 129.52 (CH, C4’’), 130.4 (CH, C5’’), 158.9 (C, C3’), 167.9 (C, C5’). HRMS-ESI+ *m/z* [M+H]^+^ calcd for [C_22_H_29_N_2_OS]^+^: 369.1995, found: 369.1997. Purity by HPLC at 254 nm: 97.65%.

### 4-((5-Bromothiophen-2-yl)methyl)-4-azatetracyclo[5.4.2.0^2,6^0^8.10^]tridecane hydrochloride (11a)

Following the general procedure B, 4-azatetracyclo[5.4.2.0^21,28^.0^37,39^]tridecane hydrochloride (120 mg, 0.56 mmol), 5-bromothiophene-2-carbaldehyde (118 mg, 0.62 mmol), NaCNBH_3_ (141 mg, 2.24 mmol) and glacial acetic acid (0.06 mL, 1.12 mmol) were mixed to obtain a yellow oil (360 mg). After column chromatography a yellow oil was obtained (176 mg, 89 % yield) that formed its hydrochloride salt as a white solid (90 mg). ^1^H NMR (400 MHz, MeOD) *δ*: 1.57 [s, 2H, 1(7)-H], 1.65 (d, *J* = 9.2 Hz, 2H, 12(13)-Ha), 2.01 (m, 2H, 12(13)-Hb), 2.13-2.23 [c.s., 4H, 9(10)-H_2_], 2.28 [broad s, 2H, 2(6)-H], 2.43 [broad s, 2H, 8(11)-H], 3.18 (t, *J* = 9.4 Hz, 2H, 3(5)-Ha], 3.62 (m, 2H, 3(5)-Hb], 4.68 (s, 2H, NH-C<u>H_2_</u>), 7.15 (d, *J* = 4.0 Hz, 1 H, 3’), 7.21 (d, *J* = 4.0 Hz, 1 H, 4’). ^13^C-NMR (100.6 MHz, MeOD) δ: 15.4 [CH_2_, C12(13)], 21.4 [CH_2_, C9(10)], 29.6 [CH, C1(7)], 38.0 [CH, C8(11)], 39.2 [CH, C2(6)], 51.7 (CH_2_, NH-<u>C</u>H_2_), 56.3 [CH_2_, C3(5)], 116.3 (C, C2’), 132.0 (CH, 3’), 133.7 (CH, C4’), 134.7 (C, C5’). HRMS-ESI+ *m/z* [M+H]^+^ calcd for [C_17_H_23_BrNS]^+^: 352.0729, found: 352.0736. Purity by HPLC at 254 nm: 99.45%.

### Biological Testing

#### Two-Electrode Voltage Clamp (TEVC) Assay

The inhibitors were tested via a TEVC assay using *X. laevis* frog oocytes microinjected with RNA expressing the M2 protein as in a previous paper. ^47^ The blocking effect of the derivatives against M2 was investigated with EP experiments using oocytes injected withA/WSN/33-M2-N31S (WT), A/WSN/33-M2-N31S-V27A (V27A), A/WSN/33-M2-N31S-L26F (L26F) and A/WSN/33 (S31N) RNAs. Mutations were generated using QuikChange site-directe mutagenesis kit (Agilent) in pGEM vector. M2 mRNAs were generated with mMessage mMachine (Thermo Fisher) using T7 promoter. The potency of the inhibitors was expressed as the inhibition percentage of the A/M2 current observed after 2 min of incubation with 100 μM of compound at pH5.5.

### Biological activity

#### Cell culture assays for anti-influenza A activity and cytotoxicity

Madin-Darby canine kidney (MDCK) cells (Cat.no. RIE 328, Friedrich-Loeffler Institute, Riems, Germany) were propagated as monolayer in Eagle’s minimum essential medium (EMEM) supplemented with 10% fetal bovine serum, 1% non-essential amino acids (NEAA), 1 mM sodium pyruvate and 2 mM L-glutamine. ^48^

The viruses used are indicated in Table 1 and including two (sub)types of influenza A virus: A/Puerto Rico/8/34 (A/H1N1) and A/Hong Kong/7/87 (A/H3N2). ^49^

Cytotoxicity and CPE inhibition studies were performed on two-day-old confluent monolayers of MDCK cells grown in 96-well plates as published. ^47^ Cytotoxicity was analyzed 72 h after compound’s addition. In CPE inhibition assay, 50 μl of a serial half-log dilution of compound in test medium (maximum concentration 100 µM) and a constant multiplicity of infection of test virus in a volume of 50 µL of the test medium were added to cells. Then, plates were incubated at 37 °C with 5% CO_2_ for 48 h. Crystal violet staining and determination of the 50% cytotoxic (CC_50_) and 50% inhibitory concentration (EC_50_) was performed as described before. ^47, 50^ At least three independent assays were conducted.

Stock solutions (10.000 µM) of all compounds were prepared in DMSO. The maximum compound concentration tested was 100 µM and consists 1 % DMS0 in cytotoxicity as well as CPE inhibition studies. Four half-log dilutions (3.16-100 µM; duplicates) were analyzed in cytotoxicity assay and at least 6 half-log dilutions (0.316-100 µM; duplicates) in CPE inhibition assays. At least two assays were conducted with inactive compounds. Active compounds were tested at least three times.

#### Anti-coronavirus Evaluation in Cell Culture

HCoV-229E was purchased from ATCC (VR-740) and expanded in human embryonic lung fibroblast cells (HEL; ATCC CCL-137). The titers of virus stocks were determined in HEL cells and expressed as TCID50 (50% tissue culture infective dose). ^51^ The cytopathic effect (CPE) reduction assay was performed in 96-well plates containing confluent HEL cell cultures, as previously described. ^52^ Serial compound dilutions were added together with HCoV-229E at an MOI of 100. In parallel, the compounds were added to a mock-infected plate to assess cytotoxicity. Besides the test compounds, one reference was included GS-441524 (the nucleoside form of remdesivir; from Carbosynth). After 5 days incubation at 35 °C, microscopy was performed to score virus-induced CPE. To next perform the colorimetric MTS cell viability assay, the reagent (CellTiter 96 AQueous MTS Reagent from Promega) was added to the wells, and 24 h later, absorbance at 490 nm was measured in a plate reader. Antiviral activity was calculated from three independent experiments and expressed as EC50 or concentration showing 50% efficacy in the MTS or microscopic assay (see ref ^53^ for calculation details). Cytotoxicity was expressed as 50% cytotoxic concentration (CC50) in the MTS assay.

### DMPK assays

#### Microsomal stability

The rat and mice pooled microsomes, with a protein content of 20 mg mL^‒1^, were purchased from Tebu-Xenotech. The compounds were incubated in a 96-well microplate at 37 °C with the microsomes in 50 mM phosphate buffer (pH = 7.4) containing as cofactors 30 mM MgCl_2_, 10 mM NADP, 100 mM glucose-6-phosphate and 40 U mL^‒1^ glucose-6-phosphate dehydrogenase, using the following volumes: 300 mL PBS, 163 mL cofactors, 5 mL test compounds (at 5 mM) and 30 mL microsomes. Samples (75 μL) were taken from each well at 0, 10, 20, 40 and 60 min and transferred to a microplate. Acetonitrile (75 μL) and internal standard (rolipram) were then added for inactivating the microsomes, and water with 0.5% formic acid (30 μL) was subsequently added for improving the chromatographic conditions, keeping the mixtures at 4 °C. The plate was centrifuged at 46,000 g for 30 min at 15 °C and supernatants were taken and analyzed in a UPLC-MS/MS system (UPLC QSM Waters Acquity) using reverse phase Acquity BEH C18 1.7 μm (2.1 mm × 50 mm, Waters) as the stationary phase, 0.1% formic acid in water (A) / 0.1% formic acid in acetonitrile (B) as the mobile phase, a flow of 0.6 mL min^‒1^. The gradient was 95% A and 5% B at 0, 0.1 min; 100% at 1, 2.1 min and 95% A 5% B at 2.5. The metabolic stability of the compounds was calculated from the logarithm of the remaining compound at each of the time points studied.

#### Permeability

The Caco-2 cells were cultured to confluency, trypsinized and seeded onto a filter transwell inserted at a density of ∼10,000 cells/well in DMEM cell culture medium. Confluent Caco-2 cells were sub-cultured at passages 58-62 and grown in a humidified atmosphere of 5% CO_2_ at 37°C. Following an overnight attachment period (24 h after seeding), the cell medium was replaced with fresh medium in both the apical and basolateral compartments every other day. The cell monolayers were used for transport studies 21 days post seeding. The monolayer integrity was checked by measuring the transepithelial electrical resistance (TEER) obtaining values ≥ 500 Ω/cm^2^. On the day of the study, after the TEER measurement, the medium was removed and the cells were washed twice with pre-warmed (37°C) Hank’s Balanced Salt Solution (HBSS) buffer to remove traces of medium. Stock solutions were made in dimethyl sulfoxide (DMSO), and further diluted in HBSS (final DMSO concentration 1%). Each compound and reference compounds (Colchicine, E3S) were all tested at a final concentration of 10 μM. For A → B directional transport, the donor working solution was added to the apical (A) compartment and the transport media as receiver working solution was added to the basolateral (B) compartment. For B → A directional transport, the donor working was added to the basolateral (B) compartment and transport media as receiver working solution was added to the apical (A) compartment. The cells were incubated at 37°C for 2 hours with gentle stirring.

At the end of the incubation, samples were taken from both donor and receiver compartments and transferred into 384-well plates and analyzed by UPLC-MS/MS. The detection was performed using an ACQUITY UPLC /Xevo TQD System. After the assay, Lucifer yellow was used to further validate the cell monolayer integrity, cells were incubated with LY 10μM in HBSS for 1hour at 37°C, obtaining permeability (P_app_) values for LY of ≤ 10 nm/s confirming the well-established Caco-2 monolayer.

#### hERG inhibition assay

The assay was carried out at a CHO cell line transfected with the hERG potassium channel. 72h before the assay, 2500 cells were seeded on a 384 well black plate (Greiner 781091). Cell line were maintained at 37°C in a 5% CO_2_ atmosphere for 24h and at 30°C in a 5% CO_2_ atmosphere for 48h plus. hERG activity was measured by using the FluxorTM Potassium Ion Chanel Assay Kit (Thermo Fisher F10016). Medium was replaced for 20μl Loading Buffer and the cells were incubated for 60 minutes at RT, protected from direct light. After incubation, Loading Buffer was replaced for Assay buffer and the compounds were incubated for 30 minutes at RT. 5μl of Stimulus Buffer was added to each well and the fluorescence was read (λ_ex_=490 nm, λ_em_=525nm) using imaging plate reader system (FDSS7000EX, Hamamatsu^®^) every second after the establishment of a baseline line.

#### Cytochrome P450 inhibition assay

The objective of this study was to screen the inhibition potential of the compound using recombinant human cytochrome P450 enzymes (CYP1A2, CYP2C19, CYP3A4 (BFC) and probe substrates with fluorescent detection. Incubations were conducted in a 200 µL volume in 96-well microtiter plates (COSTAR 3915). The addition of the mixture buffer-cofactor (KH_2_PO_4_ buffer, 1.3 mM NADP, 3.3 mM MgCl_2_, 3.3 mM glucose-6-phosphate and 0.4 U/mL glucose-6-phosphate dehydrogenase), control supersomes, standard inhibitors (furafyline, tranylzypromine, ketoconazole, sulfaphenazole and quinidine; Sigma Aldrich), and previously diluted compound to plates was carried out by a liquid handling station (Zephyr Caliper). The plate was then preincubated at 37 °C for 5 min, and the reaction was initiated by the addition of prewarmed enzyme/substrate (E/S) mix. The E/S mix contained buffer (KH_2_PO_4_), c-DNA-expressed P450 in insect cell microsomes, substrate (3-cyano-7-ethoxycoumarin, for CYP1A2 and CYP2C19; and dibenzylfluorescein for CYP3A4) in a reaction volume of 200 µL. Reactions were terminated after various times (a specific time for each cytochrome) by addition of STOP solution (ACN/TrisHCl 0.5 M 80:20 or 2 N NaOH). Fluorescence per well was measured using a fluorescence plate reader (Tecan Infinity M1000 pro) and percentage of inhibition was calculated.

### Computational methods

#### Protein preparation

To model M2(22-46) S31N proton channel we used the structure with PDB ID 2KQT^54^, which corresponds to the transmembrane domain of the WT channel (residues 22 – 46), in complex with amantadine. The preparation was performed with the protein preparation wizard^55^ panel within the Maestro interface (Schrödinger Release 2022-1: Maestro, Schrödinger, LLC, New York, NY, 2021). The pH was set to 7 ± 0.5, missing hydrogen atoms were added, Ν- and C-termini of the M2 model systems were capped by acetyl and methylamino groups, respectively. The H37 residues were protonated at the Ν_ε_ site according to experimental data. ^56–59^ All atoms of the protein complex were minimized using the OPLS2005 force field. ^60–62^ The structure was aligned with the appropriate coordinates from the OPM Server, ^63^ so the protein would have the correct orientation as regards membrane. Finally, by using the build panel within Maestro we mutated serine-31 to asparagine, amantadine was deleted, and performed one more minimization step with distance-dependent dielectric constant 4.0 using the conjugate gradient method until the root-mean-square (RMSD) of potential energy reached 2.4 × 10^−5^ kcal mol^−1^ Å^−1^ by means of Maestro/ Macromodel. ^64^

#### Docking calculations

Structures of the **6c** and **10c** were sketched with the Maestro program and model-built with the Schrödinger 2021-2 platform (Schrödinger Release 2021-2: Maestro, Schrödinger, LLC, New York, NY, 2021; Maestro-Desmond Interoperability Tools, Schrödinger, New York, NY, 2021). The models were minimized using the conjugate gradient method, MMFF94 force field ^65^, and distance-dependent dielectric constant 4.0 until a convergence threshold of 2.4 × 10^−5^ kcal mol^−1^ Å^−1^ was reached. The ligands have a charged ammonium group at pH 7.

The complexes of M2 (22-46) – **6c** and M2 (22-46) – **10c** were generated using docking calculations. A template structure was generated based on the M2 S31N (19-69) in complex with M2WJ332 (PDB ID 2LY0^66^). The M2TM S31N generated as described previously was superimposed with (PDB ID 2LY0^66^) to obtain a complex S31N M2TM - M2WJ332 which was used as template structure for docking calculations of conjugates inside S31N M2TM. **6c** and **10c** were docked inside S31N M2TM using the Induced Fit ^67–69^ standard protocol within Maestro (Schrödinger Release 2022-1: Maestro, Schrödinger, LLC, New York, NY, 2021). The access of M2WJ332 was docked inside S31N M2TM leading to a docking pose which was almost identical with the template structure, having an RMSD of the ligand ∼ 1 Å.

#### MD simulations setup

The apoprotein or its complexes **6c** and M2CD from docking calculations were aligned with the membrane using the Orientations of Proteins in Membranes (OPM) Server. ^70^ The insertion of the OPM – protein complexes into the bilayer was performed with the Desmond System Builder utility of Maestro (Schrödinger Release 2021-2: Desmond Molecular Dynamics System, D. E. Shaw Research, New York, NY, 2021. Maestro-Desmond Interoperability Tools, Schrödinger, New York, NY, 2021-2).

To generate the simulation box of each of the two systems the CHARMM – GUI ^71^ website was used (Membrane builder^71–75)^. Protein–ligand complexes were inserted in pre-equilibrated hydrated bilayers of POPC lipids expanding from the furthermost vertex of the protein to the simulation box edge 12 Å in the *x,y*-plane and 12 Å in *z*-axis. For the ligand parameters we used antechamber and GAFF2, ^76,77^ for the protein, for lipids and ions we used the CHARMM36m^78–81^ and for the water TIP3P. Intermolecular interactions were calculated using the CHARMM36m^78–81^. In the resulting system the bilayer consists of approximately 160 1-palmitoyl-2-oleoyl-sn-glycero-3-phosphocholine (POPC) lipids and 8300 waters. Sodium and chloride ions were added randomly to reach a concentration of 0.150 M. The total system contains 50000 atoms. The simulation box had dimensions of ∼ 75×75×80 Å^3^ and periodic boundary conditions were applied.

#### MD simulations protocol

The MD simulations were performed with the Gromacs 2023 ^82^ Before the MD simulations the systems were minimized and equilibrated. For the minimization protocol, we used the steep integrator with 5,000 steps with an energy tolerance of 1000 kJ/mol and constrains on hydrogen bonds 4000, 2000, 1000 kJ/mol Å^2^ on protein backbone, sidechain, and lipid atoms, respectively. This step was performed on a CPU. The equilibration protocol used MD simulations integrator and consisted by 6 equilibration steps, with restraints in hydrogen bonds gradually decreasing. The simulation time was in each step: 125, 125, 125, 250, 250 and 250 ps. The first two equilibration steps were performed in NVT ensemble, and the rest of the steps in NPT ensemble. The equilibration period followed an unrestrained 500ns-MD simulation production at temperature of 310 K ensure that the membrane state was above the melting temperature for POPC lipids.^83^ For the production the Nose – Hoover thermostat^84^ and Parinello-Rahman barostat^85^ were used. During this period the system the RMSD of the Cα carbons reached a plateau and the system was considered that reached equilibration. Two replicas, each with randomized velocities, were performed for each MD simulation. The production runs were performed with GPU acceleration.

#### Analysis of the MD simulations

The trajectories were analyzed via VMD^86^ and Gromacs^82^ tools. The water density was calculated as a linear combination of Gaussians by using the GROmaρs tool.^87^ The plots were generated using python3.10 and modules pandas^88^, matplotlib^89^ and Gromacs^81^ rms command.

## Supporting Information

Figures S1-S3; Figure S1 includes ^1^H and ^13^C NMR data; Figure 2 HPLC plots; Figure S3 the RMSD plot of the apo protein M2(22-46) S31N from the 500ns-MD simulation.

## Acknowledgements

This study was supported by funding from Fundació La Marató de TV3 (Nos. 201832 to L.N. and S.V.), and Agència de Gestió d’Ajuts Universitaris i de Recerca (grants 2021SGR00357).

## Author Information

S.V., A.L.T., and A.K. conceived and designed the research project. A.L.T. and S.V. supervised the overall research. A.L.T. and R.L. synthesized, purified, and characterized the compounds. C.M. and J.W. performed the EP measurements. K.G. carried out the molecular simulations in A.K.’s laboratory. L.N. conducted the antiviral assays. DMPK studies were performed by J.M.B., C.V., and M.I.L. A.L.T. drafted the manuscript, and all authors reviewed and approved the final version.

