## Supporting information for "Exploring polycyclic scaffolds as adamantane replacements in M2 channel inhibitors of Influenza A virus"

### CONTENTS

Table S1 S3

Table S2 S3

$^{1}H$ and $^{13}C$ NMR spectra of the new compounds S4

HPLC traces of the new compounds S17

RMSD(Ca) of M2 S31N from 500ns-MD simulations S21

**Table S1.** Antiviral activity in influenza A virus – infected MDCK cells.[a]

| | Antiviral $EC_{50}$[b,c] (µM) | | | | Cytotoxicity (µM) at 72 h |
|---|---|---|---|---|---|
| | A/H1N1 | | A/H3N2 | | |
| | CPE | MTS | CPE | MTS | MCC[d] |
| **4** | > 100 | > 100 | > 100 | > 100 | > 500 |
| **6** | 195 | 198 | 26 | 26 | > 500 |
| **6c** | | | >200 | >200 | 0.32 |
| **7** | 4.6 | 4.4 | > 100 | > 100 | > 100 |
| **8** | > 100 | > 100 | 6.5 ± 3 | 5.9 ± 4 | > 100 |
| **9** | > 100 | > 100 | > 100 | > 100 | > 500 |
| **10** | | | >200 | >200 | > 100 |
| **11**[e] | > 100 [f] | > 100 [f] | > 100 [f] | > 100 [f] | 4 [f] |

[a] MDCK: Madin-Darby canine kidney cells. [b] Virus strains: A/PR/8/34 (A/H1N1); A/HK/7/87 (A/H3N2). The $EC_{50}$ represents the 50% effective concentration, or compound concentration producing 50% inhibition of virus replication, as determined by microscopic scoring of the CPE at 72 h post infection, or by the MTS cell viability test. [c] All the compounds were inactive against influenza B/HK/5/72 (data not shown). [d] MCC: minimum cytotoxic concentration, or concentration producing minimal alterations in cell morphology after 72 h incubation with compound. [e] Compound **11** was active in a virus yield experiment ($EC_{99}$: < 0.08 µM; $EC_{99}$: compound concentration giving 2 $\log_{10}$ reduction in virus yield, as determined by quantifying the virus in the supernatant at 24 post infection, using an qRT-PCR based virus yield assay). [f] Rey-Carrizo M., *et al.*, *J. Med. Chem.* **2014**, 57, 5738.

**Table S2.** Antiviral activity in HCoV-229 infected HEL cells.

| | Antiviral activity (µM) | | Cytotoxicity (µM) | |
|---|---|---|---|---|
| | $EC_{50}$ (CPE)[a] | $EC_{50}$ (MTS)[b] | MCC (micr)[c] | $CC_{50}$ (MTS)[d] |
| **6a** | 11 | 10 | > 100 | > 100 |
| **10** | 4.7 | 4.5 | > 100 | > 100 |
| **GS-441524** | 3.8 | 2.8 | > 40 | > 40 |

[a] $EC_{50}$: concentration affording 50% protection against viral CPE, as determined by microscopic scoring. [b] $EC_{50}$: concentration affording 50% protection against viral CPE, as determined by the MTS cell viability assay. [c] MCC (minimal cytotoxic concentration), that is, compound concentration that produces a microscopically visible change in normal cell morphology. [d] $CC_{50}$: 50% cytotoxic concentration, as determined by MTS assay.

**2-(((2-Oxaadamantan-1-yl)amino)methyl)-5-methoxyphenol (4a)**

$^{1}$H NMR (400 MHz, $CDCl_3$)

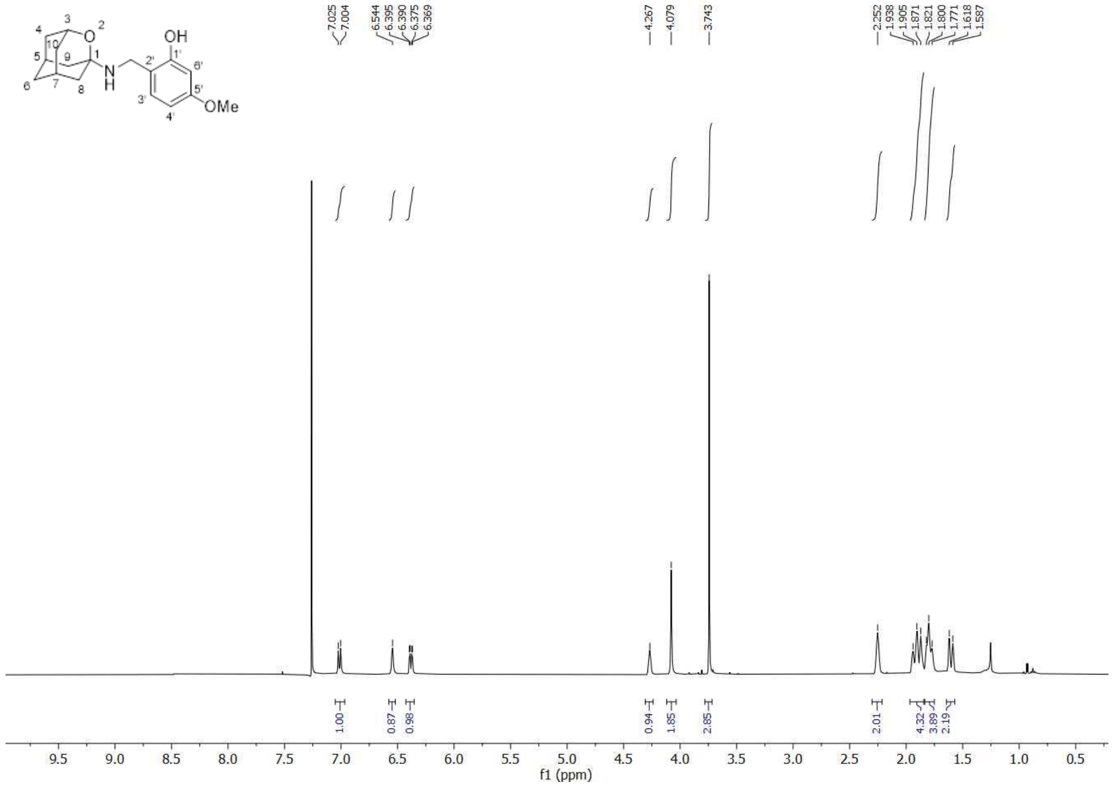


$^{13}$C-NMR (100.6 MHz, $CDCl_3$)

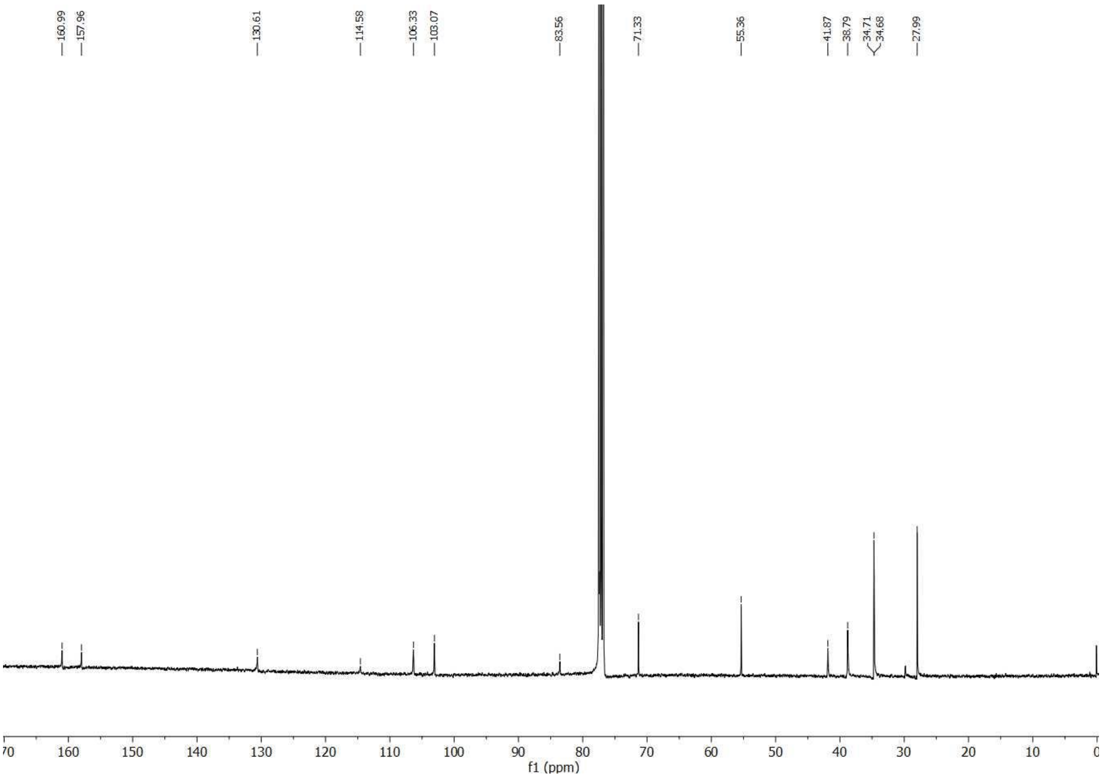

**_N_-((5-Bromothiophen-2-yl)methyl)-2-oxaadamantan-1-amine hydrochloride (4b)**

$^{1}$H NMR (400 MHz, MeOD)

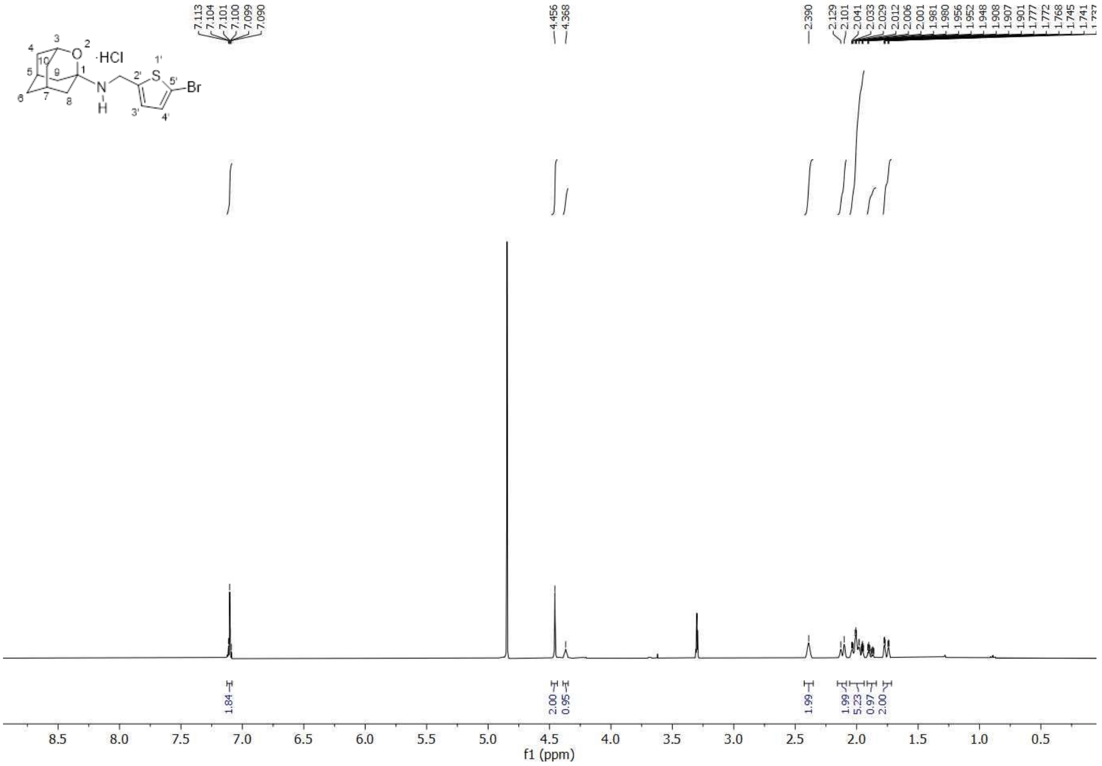


$^{13}$C-NMR (100.6 MHz, MeOD)

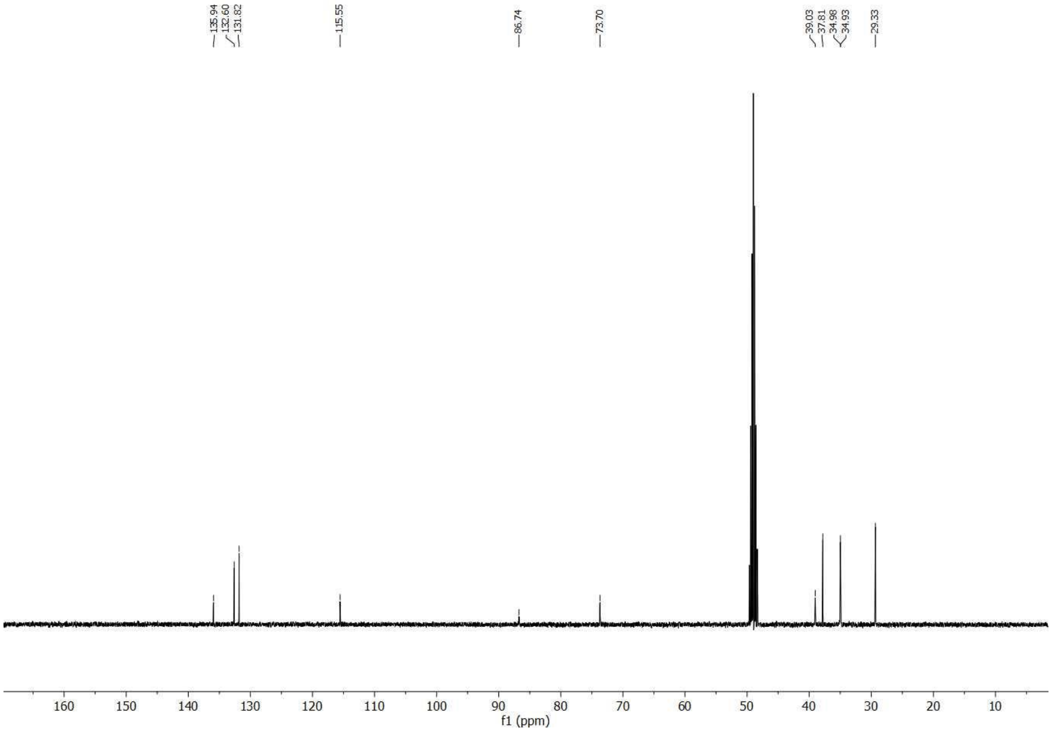

#### 2-(((2-Oxaadamantan-5-yl)amino)methyl)-5-methoxyphenol (5a)

$^{1}$H NMR (400 MHz, $CDCl_3$)

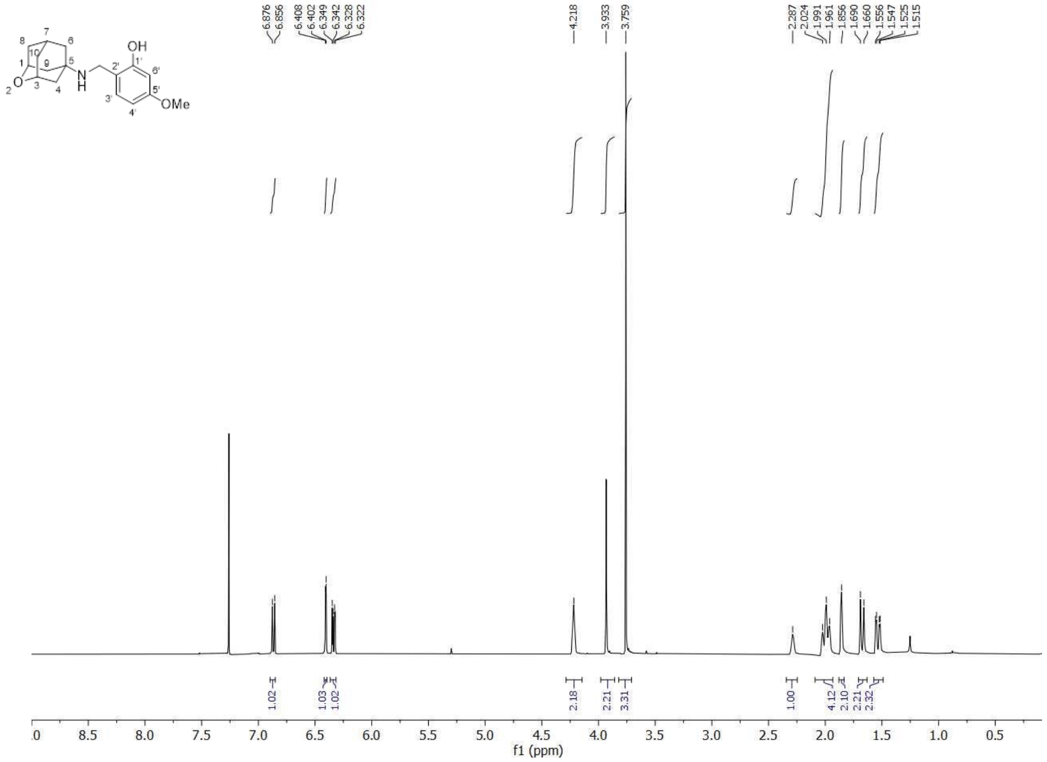


$^{13}$C-NMR (100.6 MHz, $CDCl_3$)

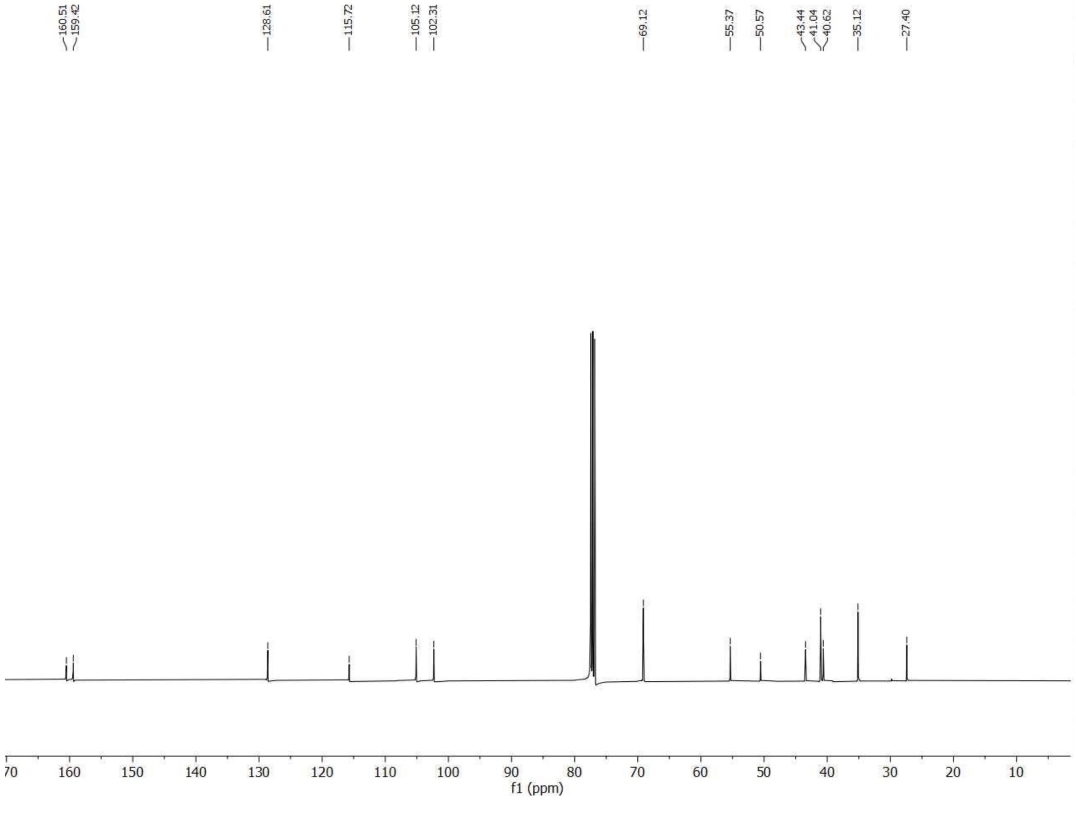

#### 2-(((Tricyclo[3.3.1.0$^{3,7}$]non-3-yl)amino)methyl)-5-methoxyphenol hydrochloride (6a)

$^{1}$H NMR (400 MHz, $CDCl_3$)

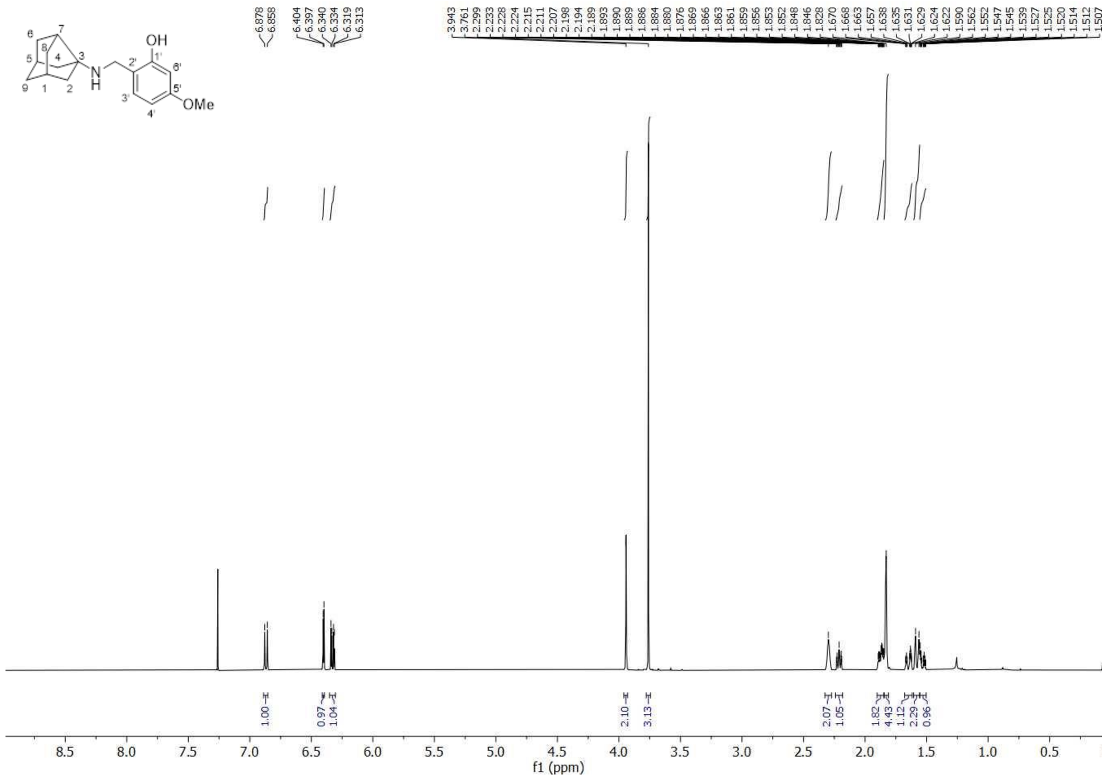


$^{13}$C-NMR (100.6 MHz, $CDCl_3$)

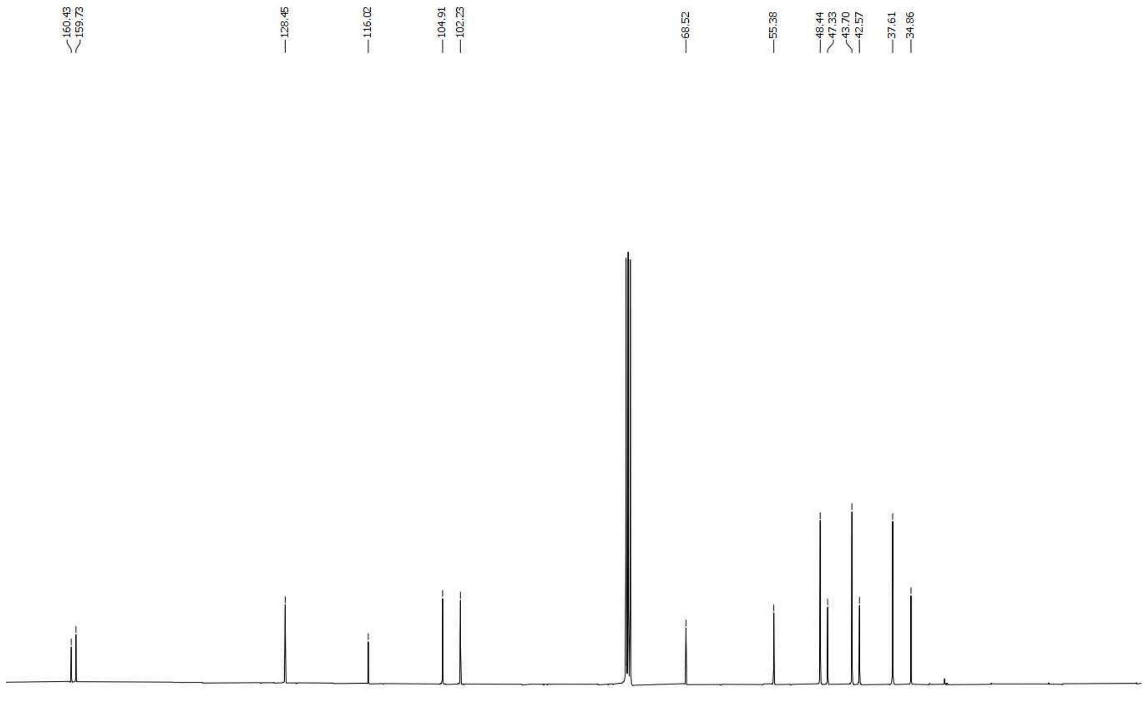

#### *N*-((5-Bromothiophen-2-yl)methyl)tricyclo[3.3.1.0$^{3,7}$]nonan-1-amine hydrochloride (6b)

$^{1}$H NMR (400 MHz, MeOD)

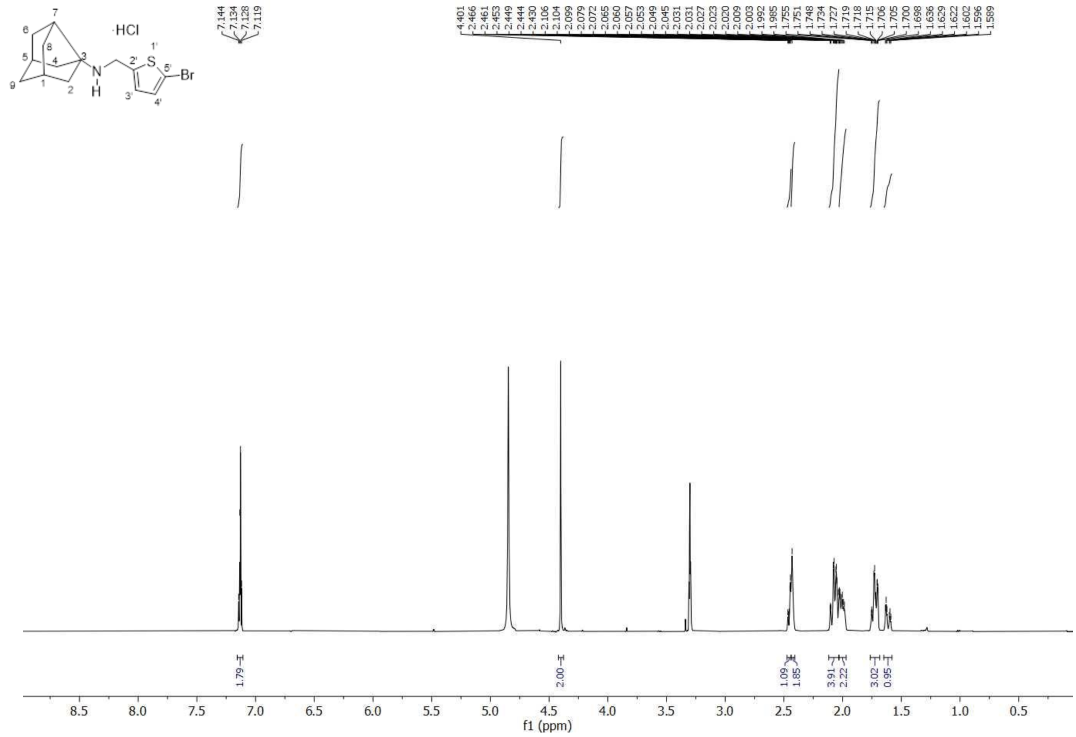

$^{13}$C-NMR (100.6 MHz, MeOD)

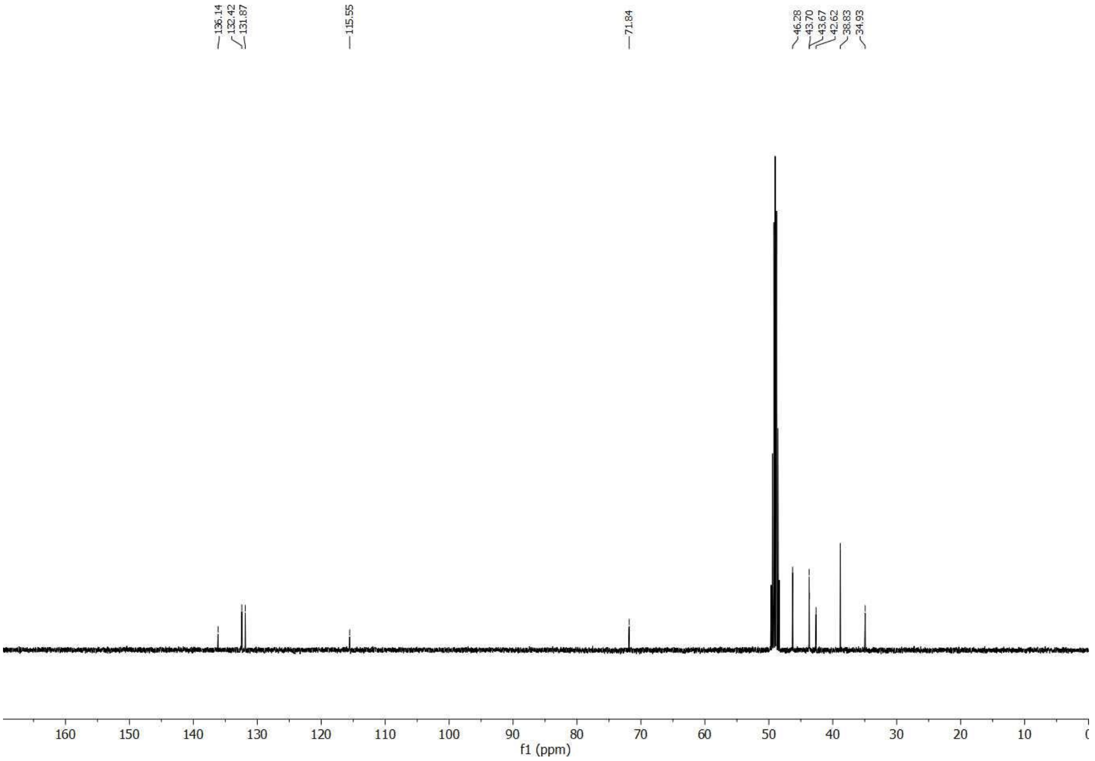

**N-((5-(Thiophen-2-yl)isoxazol-3-yl)methyl)tricyclo[3.3.1.0$^{3,7}$]nonan-1-amine hydrochloride (6c)**

$^{1}$H NMR (400 MHz, MeOD)

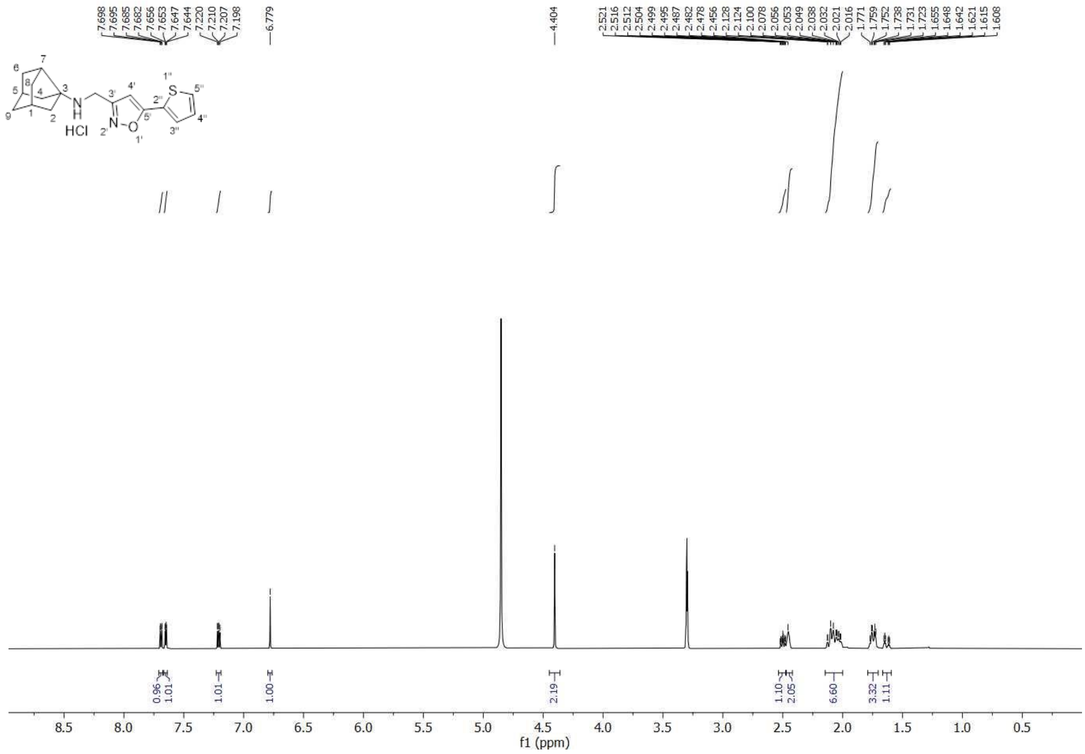


$^{13}$C-NMR (100.6 MHz, MeOD)

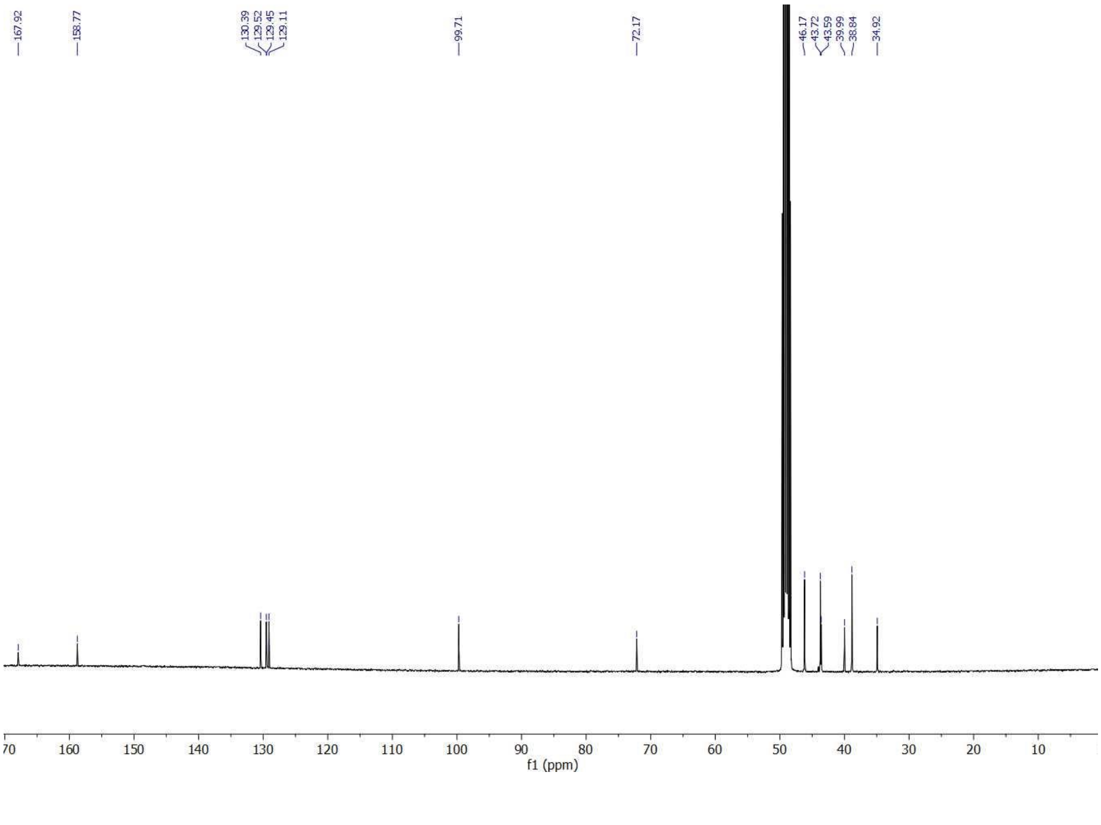

#### 5-Methoxy-2-((((3,7,9,9-tetramethyl)tricyclo[3.3.1.0$^{3,7}$]non-1-yl)amino)methyl)phenol hydrochloride (7a)

$^{1}$H NMR (400 MHz, MeOD)

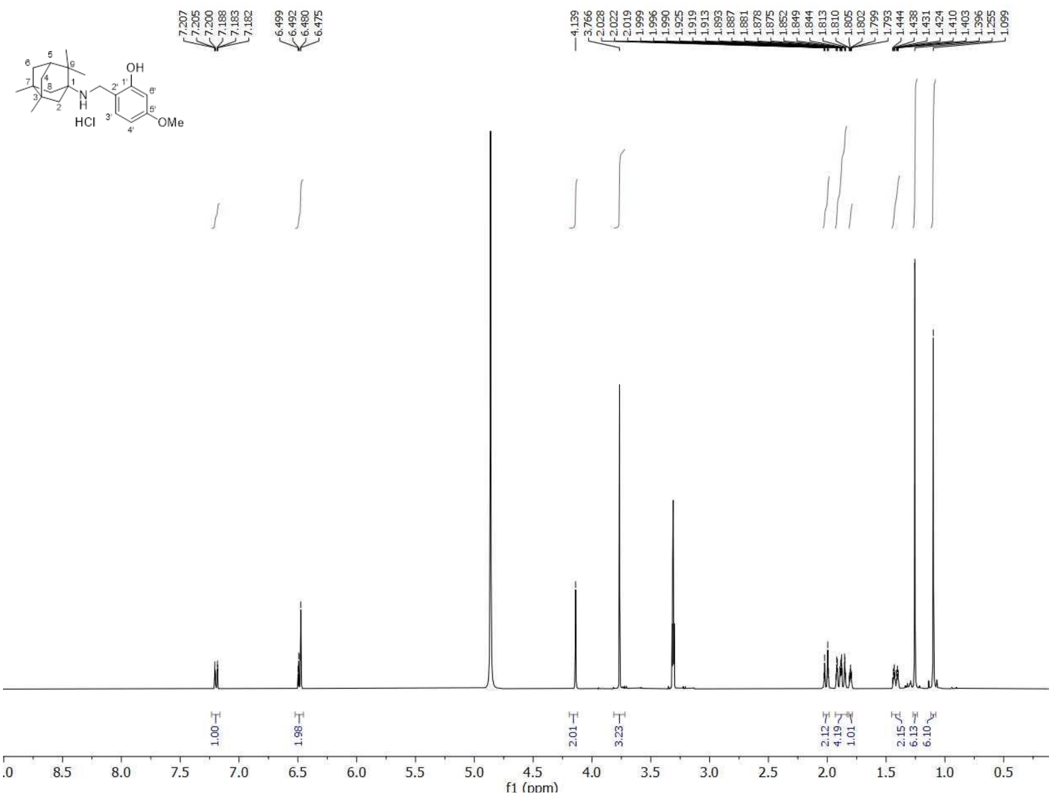


$^{13}$C-NMR (100.6 MHz, MeOD)

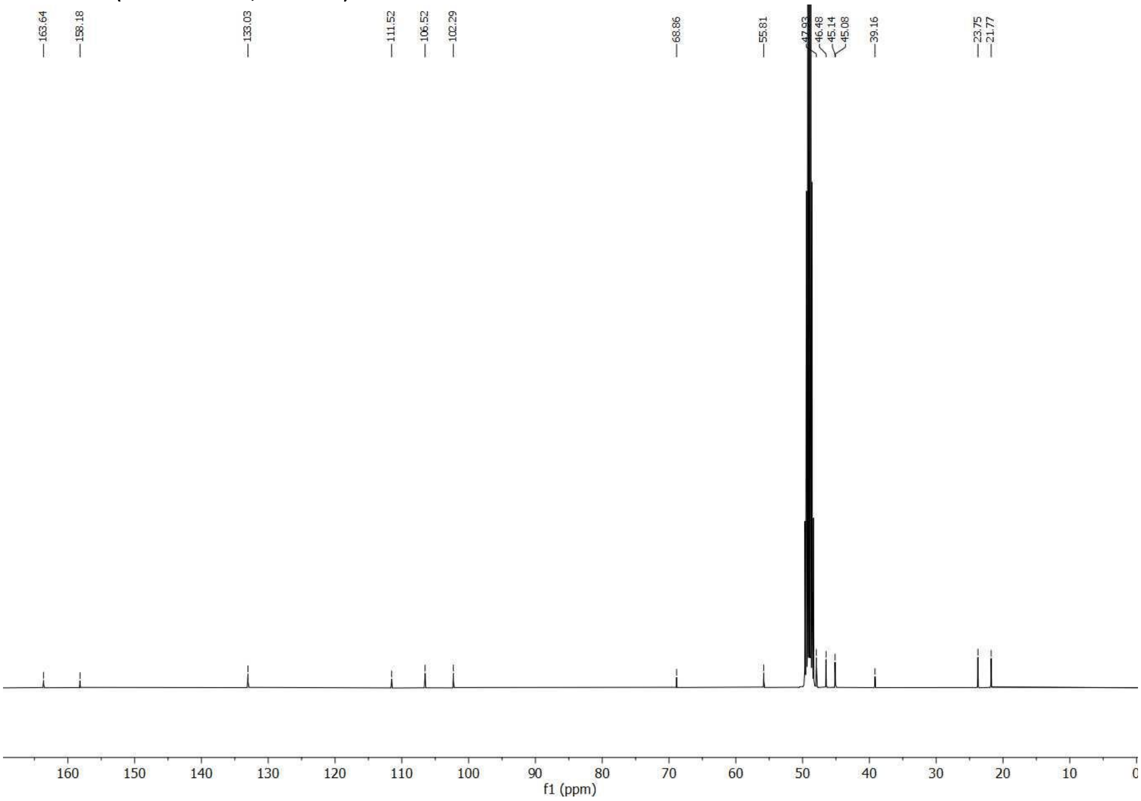

#### 2-((((3,7-Dimethyltricyclo[3.3.0.0$^{3,7}$]octan-1-yl)methyl)amino)methyl)-5-methoxyphenol hydrochloride (8a)

$^1$H NMR (400 MHz, MeOD)

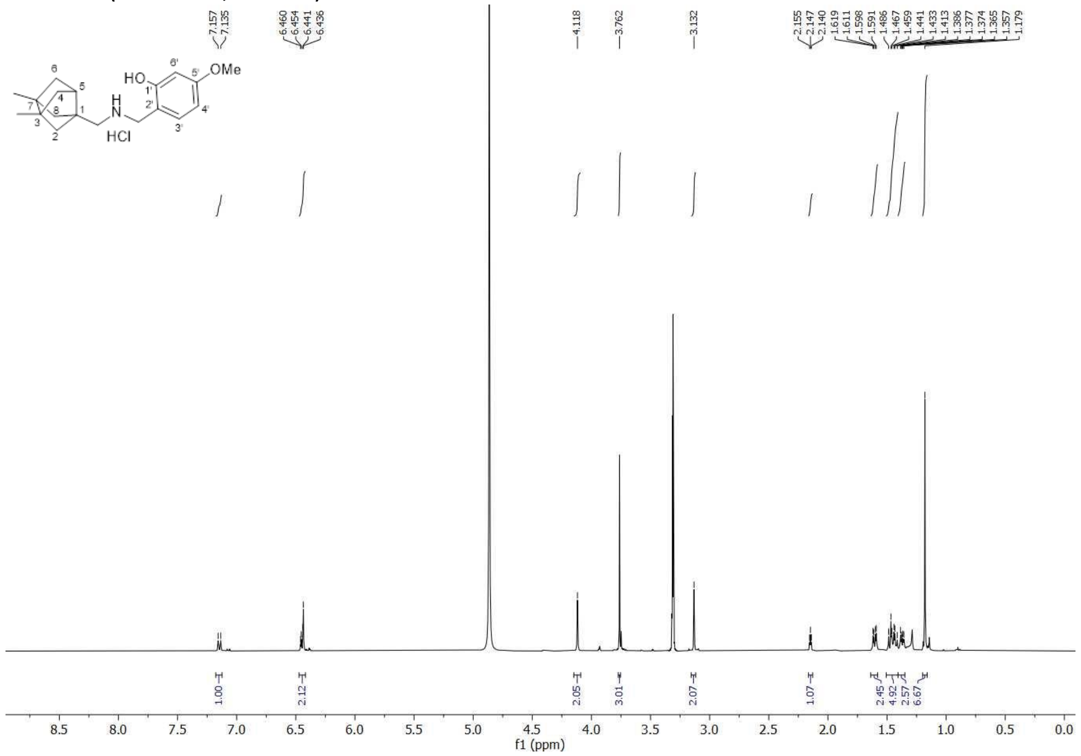


$^{13}$C-NMR (100.6 MHz, MeOD)

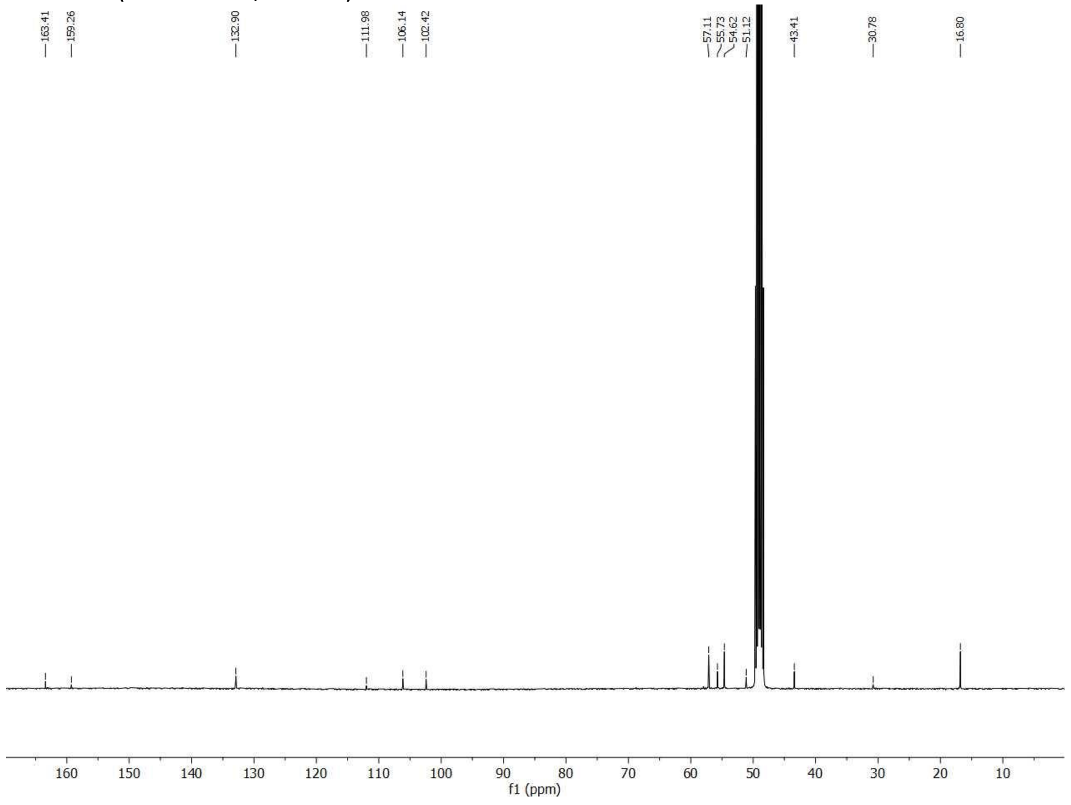

**_N_-((5-Bromothiophen-2-yl)methyl)-3,7-dimethyltricyclo[3.3.0.0$^{3,7}$]octan-1-amine hydrochloride (9a)**

$^{1}$H NMR (400 MHz, MeOD)

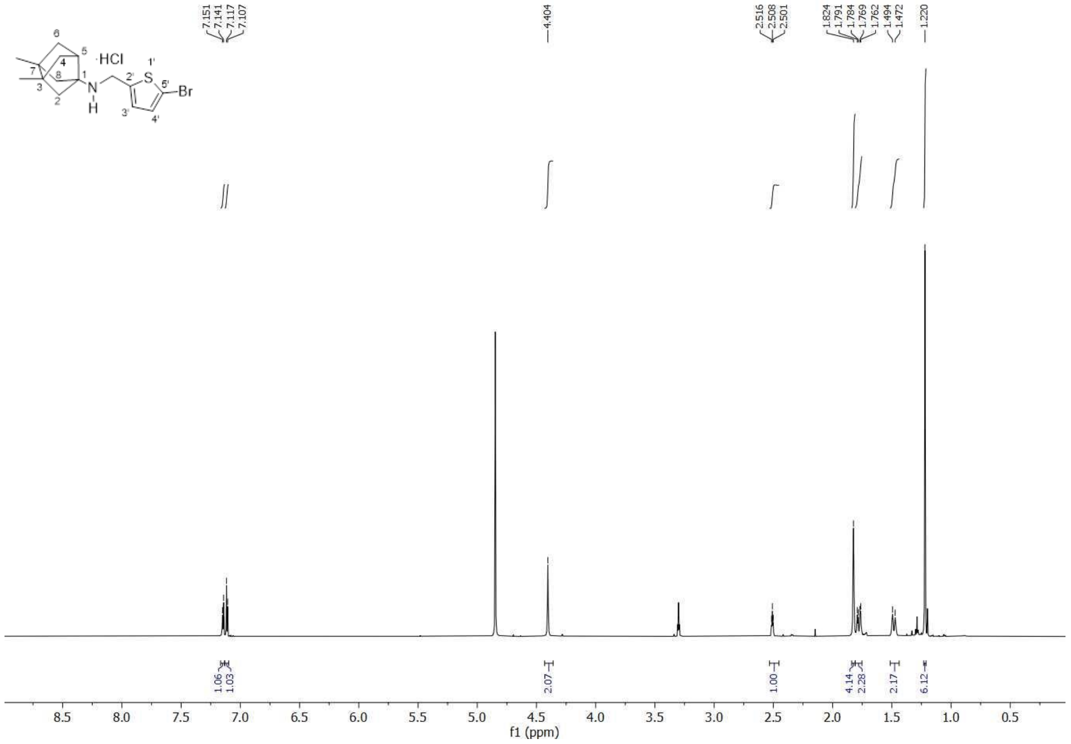


$^{13}$C-NMR (100.6 MHz, MeOD)

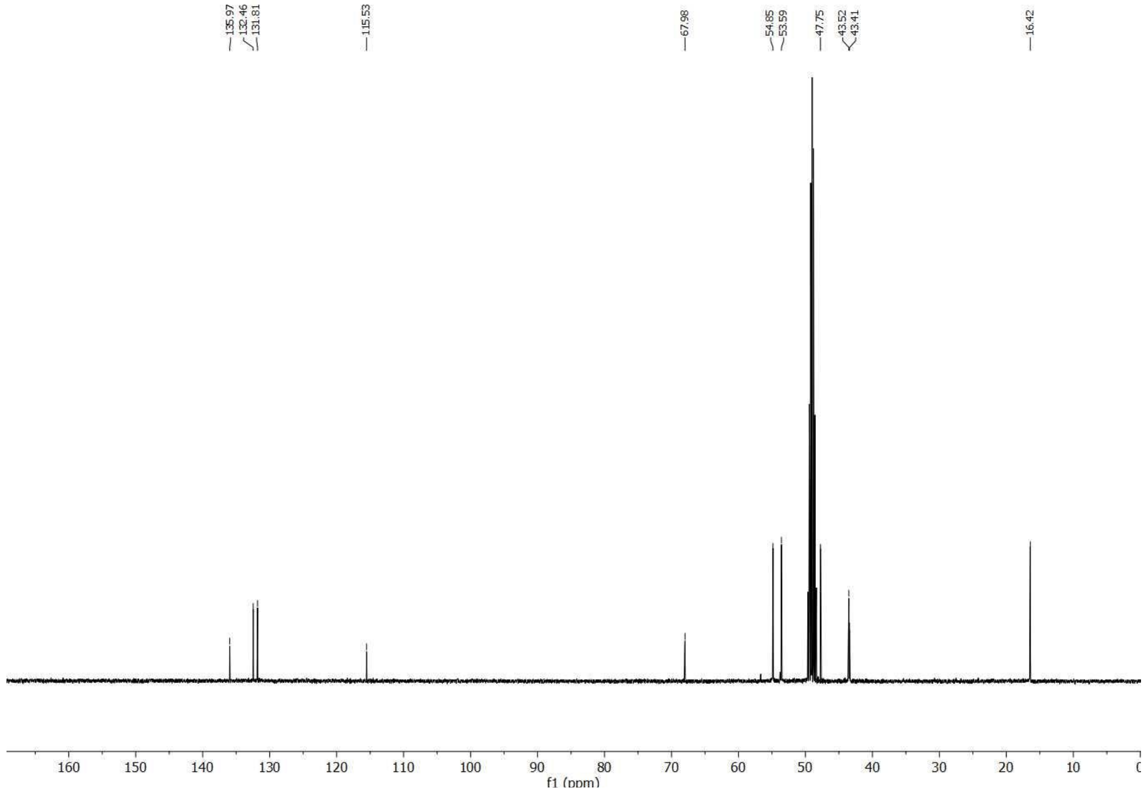

**5-Methoxy-2-(((3,4,8,9-tetramethyltetracyclo[4.4.0.0$^{3,9}$.0$^{4,8}$]decan-1-yl)amino)methyl)phenol (10a)**

$^{1}$H NMR (400 MHz, CDCl$_3$)

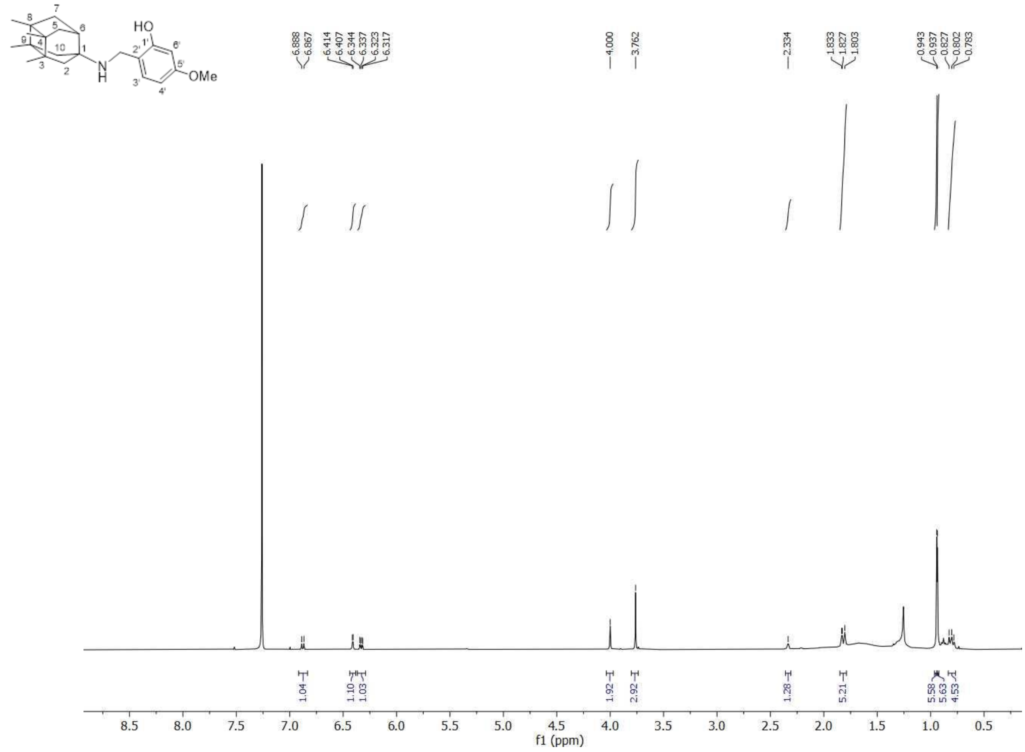


$^{13}$C-NMR (100.6 MHz, CDCl$_3$)

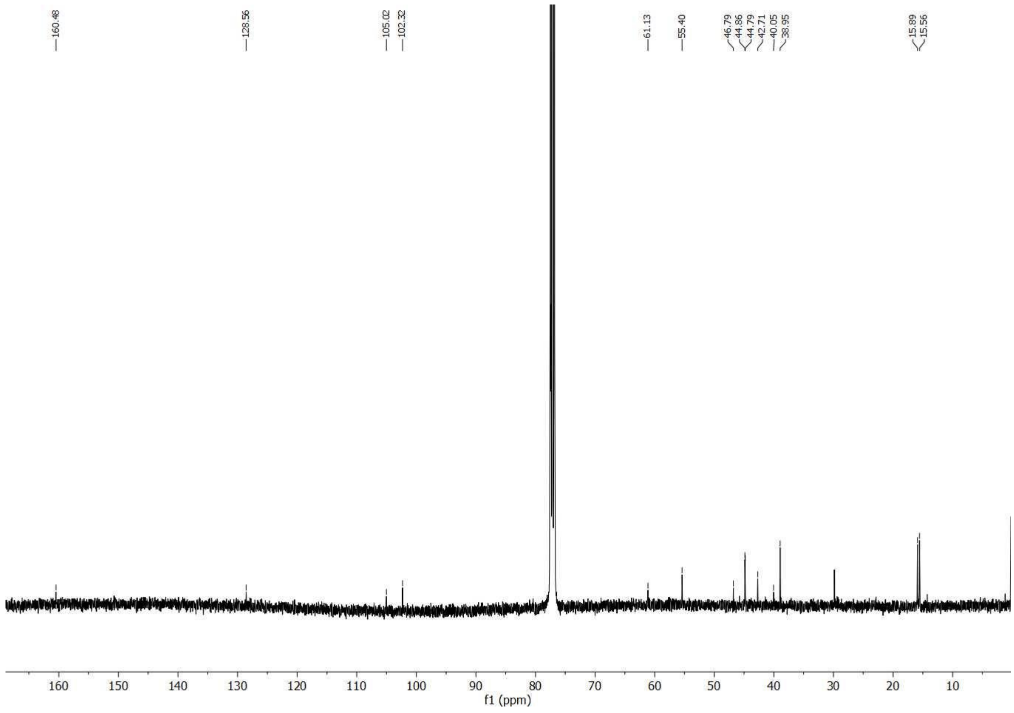

***N*-((5-Bromothiophen-2-yl)methyl)-3,4,8,9-tetramethyltetracyclo[4.4.0.0$^{3,9}$.0$^{4,8}$]decan-1-amine hydrochloride (10b)**

$^{1}$H NMR (400 MHz, MeOD)

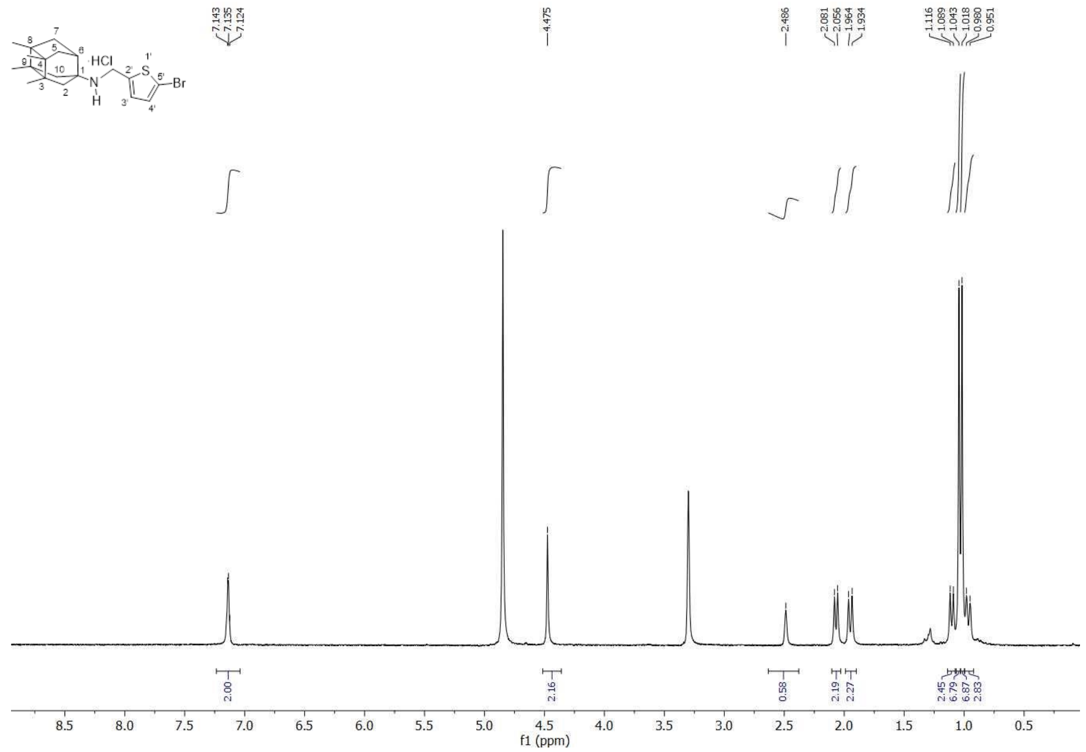

$^{13}$C-NMR (100.6 MHz, MeOD)

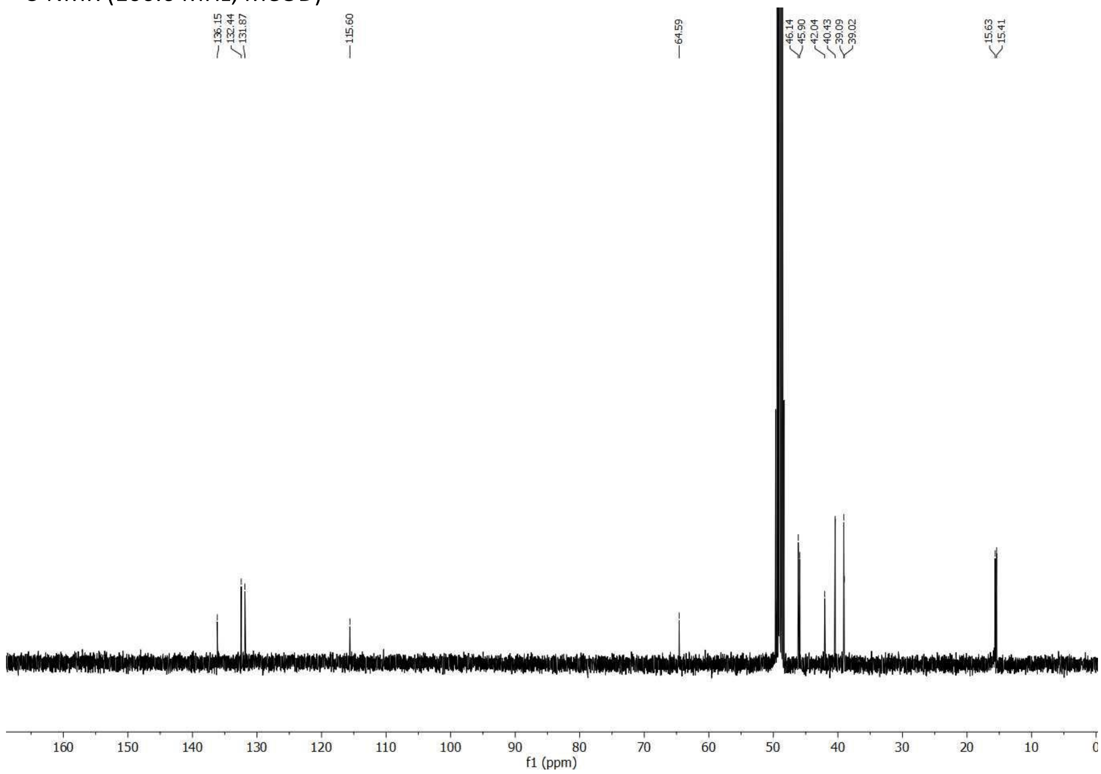

**3,4,8,9-Tetramethyl-*N*-((5-(thiophen-2-yl)isoxazol-3-yl) methyl)tetracyclo[4.4.0.0$^{3,9}$.0$^{4,8}$]decan-1-amine hydrochloride (10c)**

$^{1}$H NMR (400 MHz, MeOD)

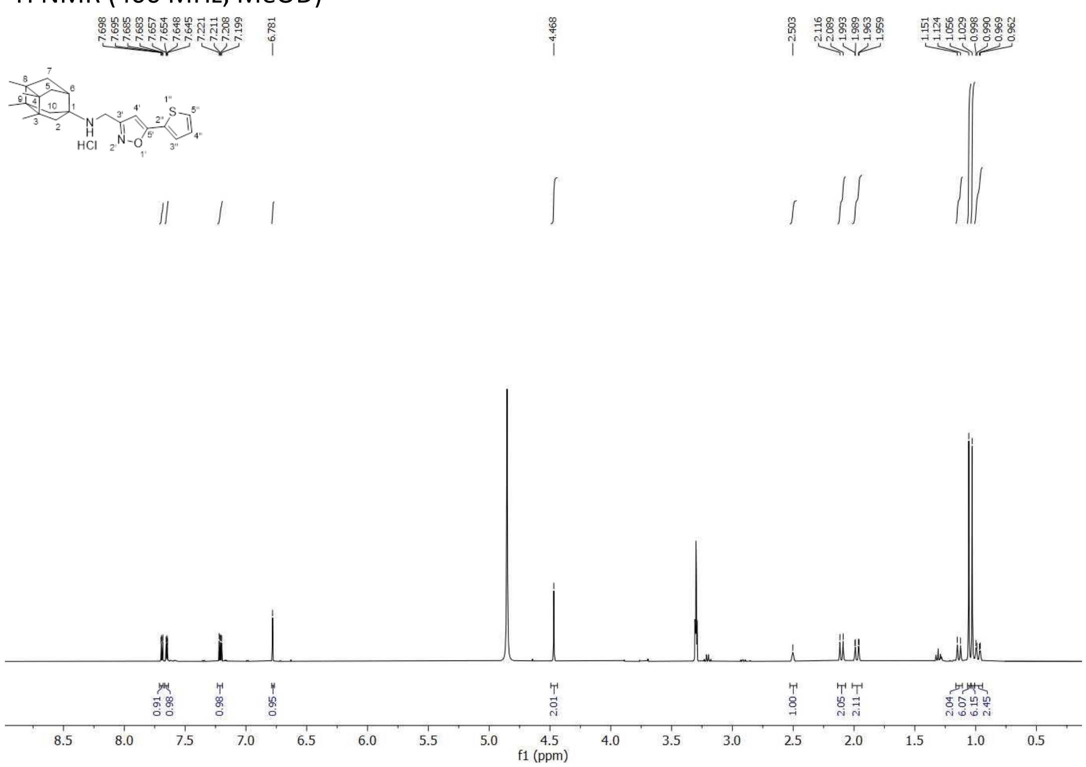


$^{13}$C-NMR (100.6 MHz, MeOD)

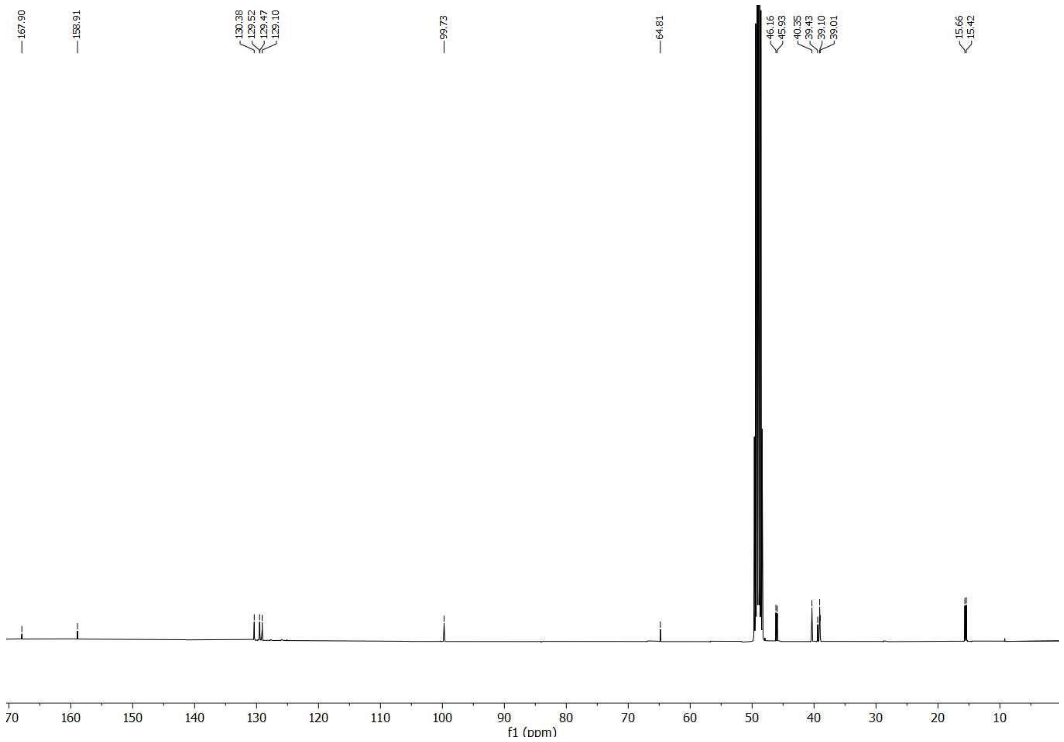

**4-((5-Bromothiophen-2-yl)methyl)-4-azatetracyclo[5.4.2.0$^{2,6}$.0$^{8.11}$]tridecane hydrochloride (11a)**

$^{1}$H NMR (400 MHz, MeOD)

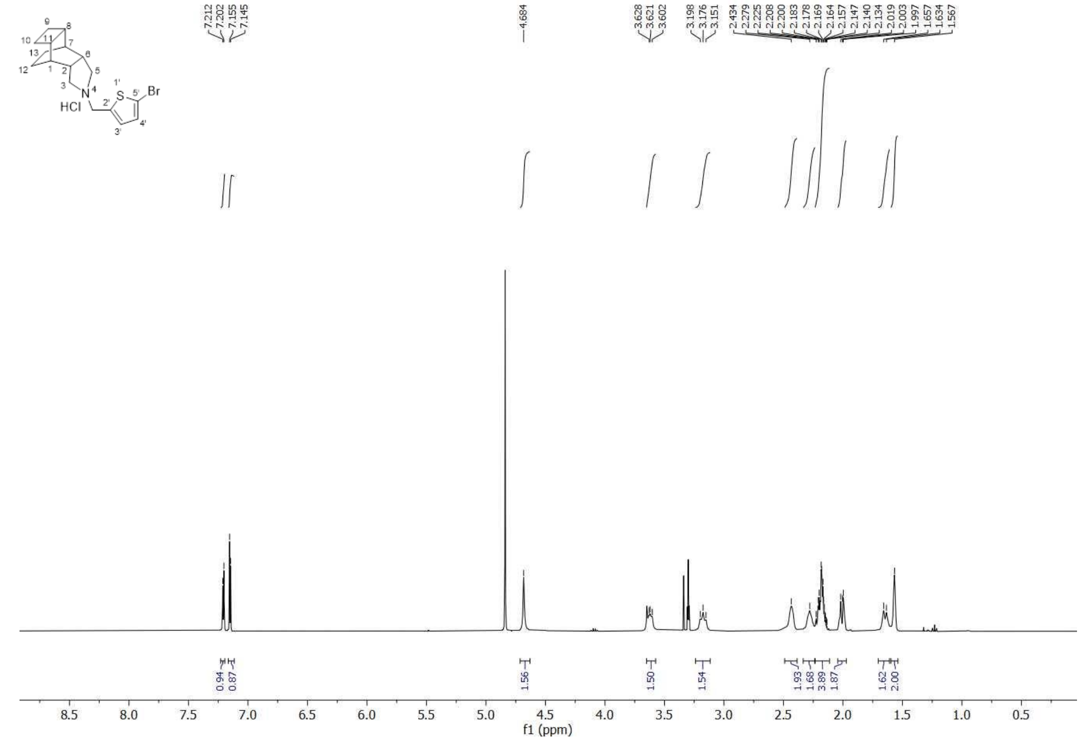


$^{13}$C-NMR (100.6 MHz, MeOD)

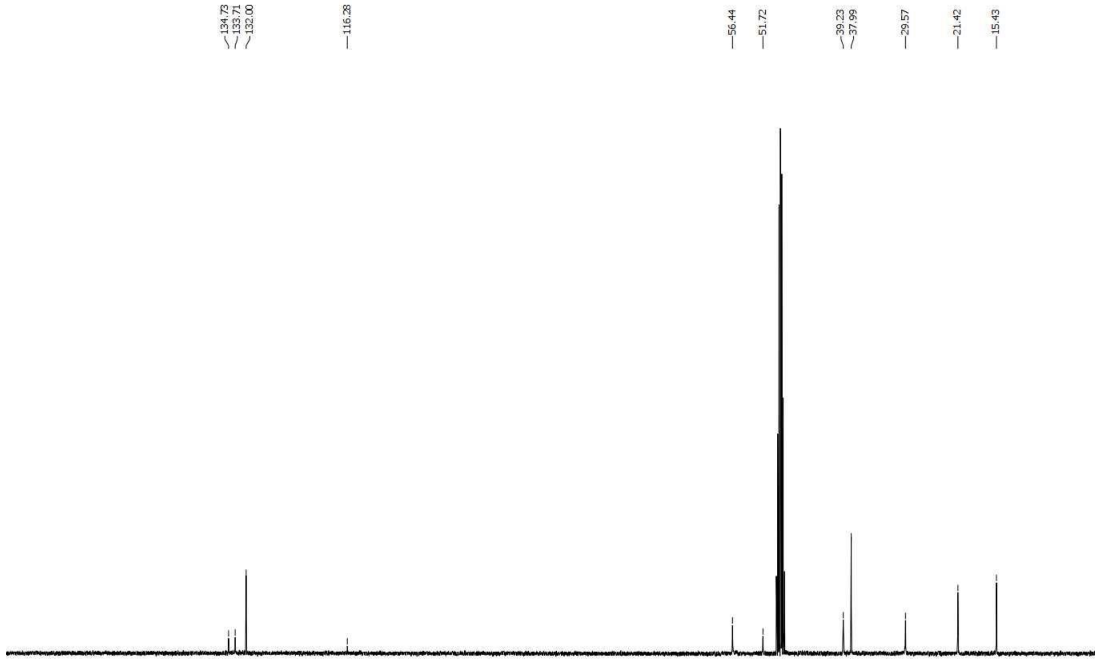


**Figure S1**. 1H and 13C NMR spectra.

#### Blank

275 nm

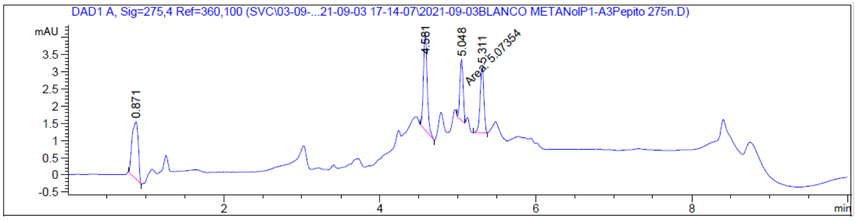


254 nm

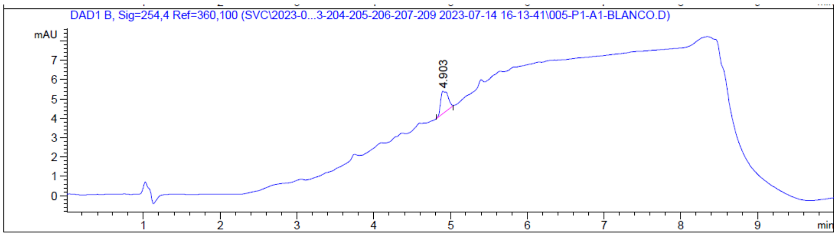


## 4a

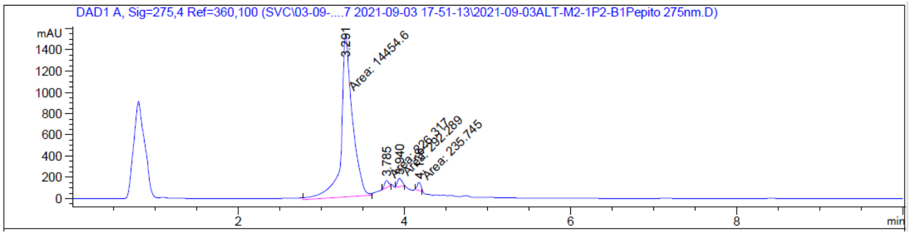


Signal 1: DAD1 A, Sig=275,4 Ref=360,100

| Peak # | RetTime [min] | Type | Width [min] | Area [mAU*s] | Height [mAU] | Area % |
|---|---|---|---|---|---|---|
| 1 | 3.291 | MM | 0.1574 | 1.44546e4 | 1530.88806 | 95.0401 |
| 2 | 3.785 | MM | 0.0562 | 226.31747 | 67.10942 | 1.4881 |
| 3 | 3.940 | MM | 0.0602 | 292.28949 | 80.92228 | 1.9218 |
| 4 | 4.178 | MM | 0.0511 | 235.74506 | 76.95282 | 1.5500 |
| Totals : | | | | 1.52089e4 | 1755.87258 | |

## 4b

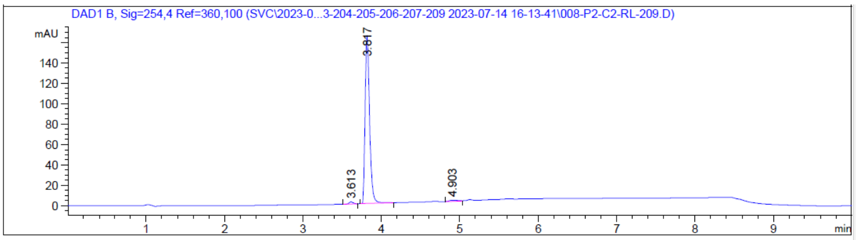


Signal 2: DAD1 B, Sig=254,4 Ref=360,100

| Peak # | RetTime [min] | Type | Width [min] | Area [mAU*s] | Height [mAU] | Area % |
|---|---|---|---|---|---|---|
| 1 | 3.613 | BB | 0.0538 | 7.00948 | 2.05650 | 1.0625 |
| 2 | 3.817 | BB | 0.0597 | 645.74219 | 164.70552 | 97.8801 |
| 3 | 4.903 | BB | 0.0861 | 6.97641 | 1.09848 | 1.0575 |
| Totals : | | | | 659.72808 | 167.86050 | |

**6a**

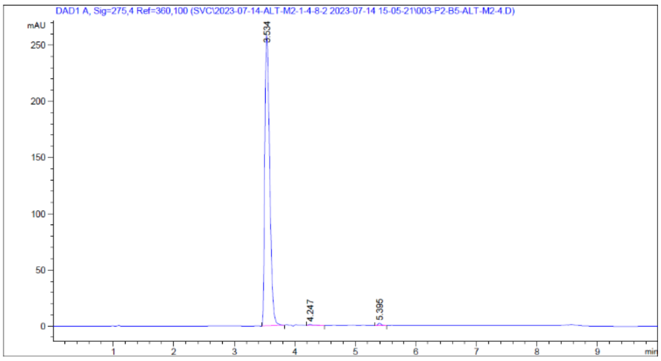


Signal 1: DAD1 A, Sig=275,4 Ref=360,100

| Peak # | RetTime [min] | Type | Width [min] | Area [mAU*s] | Height [mAU] | Area % |
|---|---|---|---|---|---|---|
| 1 | 3.534 | BB | 0.0873 | 1447.81079 | 258.81442 | 98.9865 |
| 2 | 4.247 | BB | 0.0831 | 6.87643 | 1.16181 | 0.4701 |
| 3 | 5.395 | BB | 0.0585 | 7.94670 | 2.08289 | 0.5433 |
| Totals : | | | | 1462.63392 | 262.05913 | |

**6b**

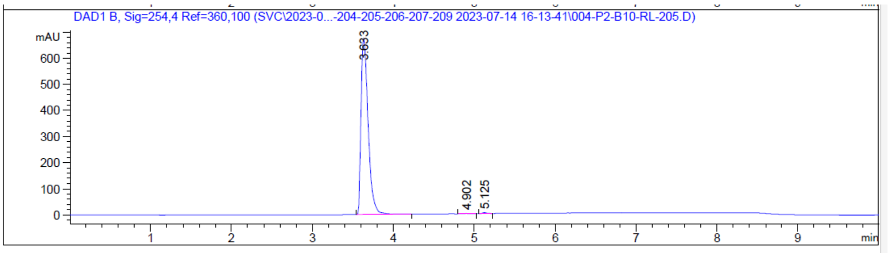


Signal 2: DAD1 B, Sig=254,4 Ref=360,100

| Peak # | RetTime [min] | Type | Width [min] | Area [mAU*s] | Height [mAU] | Area % |
|---|---|---|---|---|---|---|
| 1 | 3.633 | BB | 0.0931 | 4095.21484 | 672.04993 | 99.5014 |
| 2 | 4.902 | BB | 0.0858 | 7.10319 | 1.12273 | 0.1726 |
| 3 | 5.125 | BB | 0.0563 | 13.41812 | 3.70402 | 0.3260 |
| Totals : | | | | 4115.73615 | 676.87667 | |

**6c**

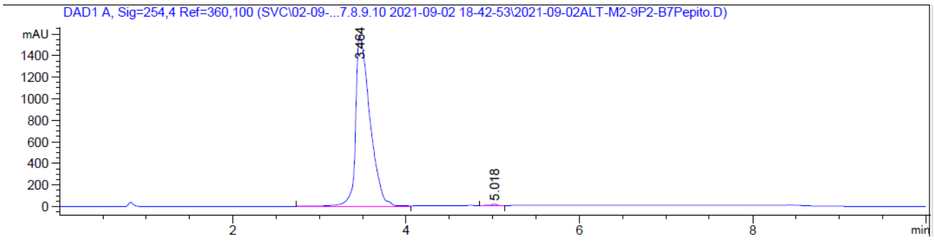


Signal 1: DAD1 A, Sig=254,4 Ref=360,100

| Peak # | RetTime [min] | Type | Width [min] | Area [mAU*s] | Height [mAU] | Area % |
|---|---|---|---|---|---|---|
| 1 | 3.464 | BB | 0.1712 | 1.81391e4 | 1588.48340 | 99.6835 |
| 2 | 5.018 | BB | 0.0613 | 57.59582 | 14.21870 | 0.3165 |
| Totals : | | | | 1.81967e4 | 1602.70210 | |

**7a**

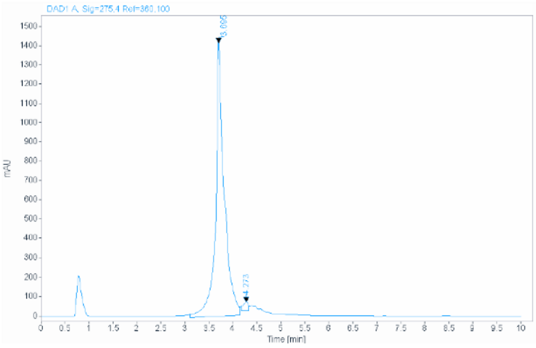

Signal: DAD1 A, Sig=275,4 Ref=360,100

| RT [min] | Type | Width [min] | Area | Height | Area% | Name |
|---|---|---|---|---|---|---|
| 3.695 | MM | 0.2084 | 17803.7422 | 1423.9562 | 98.1504 | |
| 4.273 | MM | 0.1226 | 335.4981 | 45.6213 | 1.8496 | |
| | | Sum | 18139.2403 | | | |

**8a**

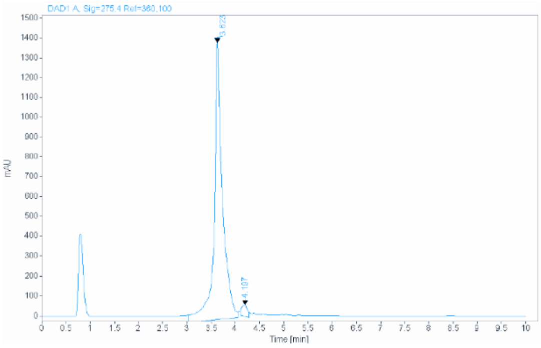

Signal: DAD1 A, Sig=275,4 Ref=360,100

| RT [min] | Type | Width [min] | Area | Height | Area% | Name |
|---|---|---|---|---|---|---|
| 3.623 | MM | 0.1970 | 16545.7695 | 1399.7637 | 97.2048 | |
| 4.197 | MM | 0.1341 | 475.7853 | 59.1475 | 2.7952 | |

**9a**

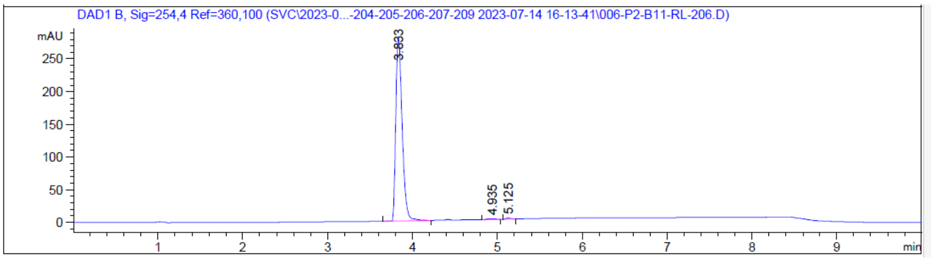


Signal 2: DAD1 B, Sig=254,4 Ref=360,100

| Peak # | RetTime [min] | Type | Width [min] | Area [mAU*s] | Height [mAU] | Area % |
|---|---|---|---|---|---|---|
| 1 | 3.833 | BB | 0.0809 | 1461.92078 | 280.28922 | 99.0313 |
| 2 | 4.935 | BB | 0.0939 | 7.28272 | 1.03692 | 0.4933 |
| 3 | 5.125 | BB | 0.0535 | 7.01710 | 1.97084 | 0.4753 |
| Totals : | | | | 1476.22059 | 283.29698 | |

**10a**

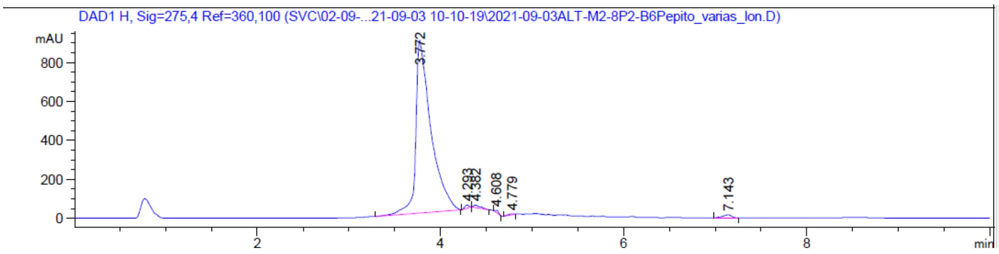

Signal 7: DAD1 H, Sig=275,4 Ref=360,100

| Peak # | RetTime [min] | Type | Width [min] | Area [mAU*s] | Height [mAU] | Area % |
|---|---|---|---|---|---|---|
| 1 | 3.772 | BB | 0.1548 | 1.02555e4 | 887.52191 | 97.3453 |
| 2 | 4.293 | BB | 0.0554 | 55.03538 | 16.27613 | 0.5224 |
| 3 | 4.382 | BB | 0.0767 | 72.86250 | 13.13648 | 0.6916 |
| 4 | 4.608 | BB | 0.0485 | 31.24779 | 10.55890 | 0.2966 |
| 5 | 4.779 | BB | 0.0931 | 22.10940 | 3.73198 | 0.2099 |
| 6 | 7.143 | BB | 0.0951 | 98.41683 | 15.29474 | 0.9342 |
| Totals : | | | | 1.05351e4 | 946.52014 | |

**10b**

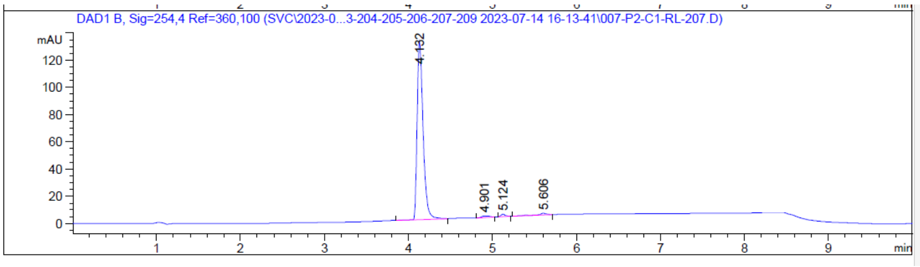


Signal 2: DAD1 B, Sig=254,4 Ref=360,100

| Peak # | RetTime [min] | Type | Width [min] | Area [mAU*s] | Height [mAU] | Area % |
|---|---|---|---|---|---|---|
| 1 | 4.132 | BB | 0.0759 | 649.10352 | 130.77872 | 96.7532 |
| 2 | 4.901 | BB | 0.0847 | 6.99991 | 1.12366 | 1.0434 |
| 3 | 5.124 | BB | 0.0558 | 6.28315 | 1.75330 | 0.9365 |
| 4 | 5.606 | BB | 0.0892 | 8.49937 | 1.31787 | 1.2669 |
| Totals : | | | | 670.88594 | 134.97354 | |

**10c**

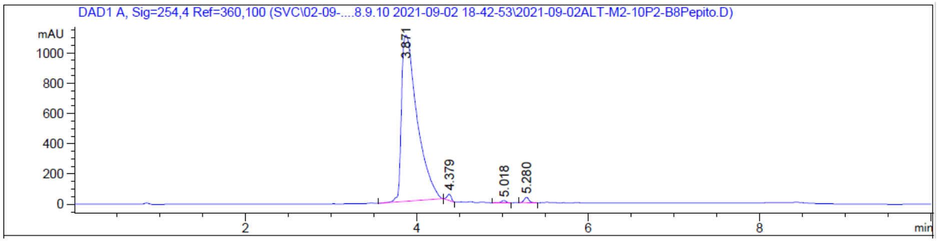


Signal 1: DAD1 A, Sig=254,4 Ref=360,100

| Peak # | RetTime [min] | Type | Width [min] | Area [mAU*s] | Height [mAU] | Area % |
|---|---|---|---|---|---|---|
| 1 | 3.871 | BB | 0.1857 | 1.39021e4 | 1098.09082 | 97.6543 |
| 2 | 4.379 | BB | 0.0510 | 138.15520 | 41.27779 | 0.9705 |
| 3 | 5.018 | BB | 0.0572 | 57.69621 | 15.59678 | 0.4053 |
| 4 | 5.280 | BB | 0.0587 | 138.08611 | 36.07040 | 0.9700 |
| Totals : | | | | 1.42360e4 | 1191.03579 | |

**11a**

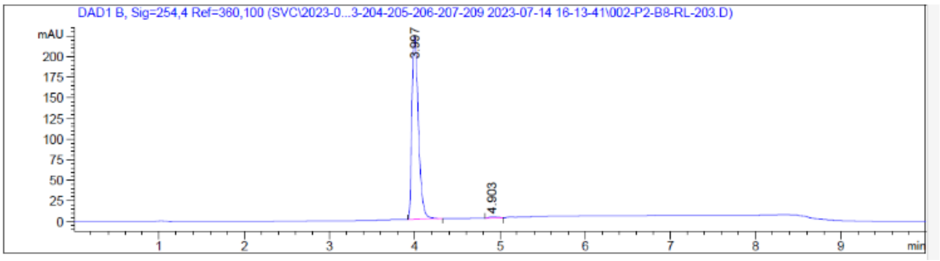


Signal 2: DAD1 B, Sig=254,4 Ref=360,100

| Peak # | RetTime [min] | Type | Width [min] | Area [mAU*s] | Height [mAU] | Area % |
|---|---|---|---|---|---|---|
| 1 | 3.997 | BB | 0.0821 | 1182.03589 | 222.10286 | 99.4526 |
| 2 | 4.903 | BB | 0.0841 | 6.50579 | 1.05274 | 0.5474 |
| Totals : | | | | 1188.54168 | 223.15560 | |

**Figure S2.** HPLC plots for the final products.

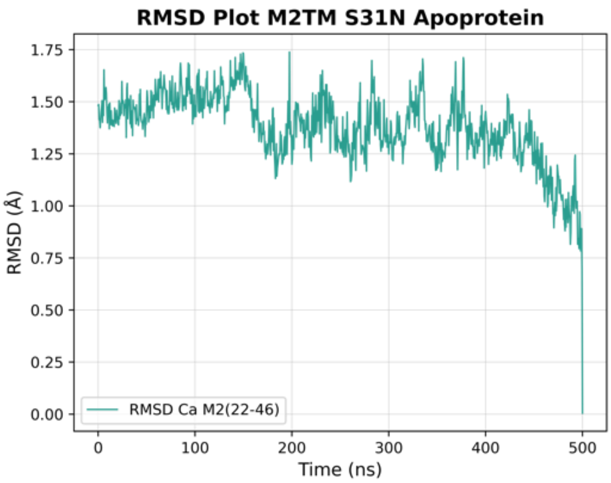


**Figure S3.** RMSD(Ca) plot from 500ns-MD simulations of apo-protein M2(22-46) S31N embedded in POPC bilayers using the CHARMM36m force field.
